# Spatially resolved molecular and cellular atlas of the mouse brain

**DOI:** 10.1101/2023.12.03.569501

**Authors:** Lei Han, Zhen Liu, Zehua Jing, Yuxuan Liu, Huizhong Chang, Junjie Lei, Yujie Peng, Kexin Wang, Yuanfang Xu, Wei Liu, Zihan Wu, Qian Li, Xiaoxue Shi, Mingyuan Zheng, He Wang, Juan Deng, Yanqing Zhong, Hailin Pan, Junkai Lin, Ruiyi Zhang, Yu Chen, Jinhua Wu, Mingrui Xu, Biyu Ren, Mengnan Cheng, Qian Yu, Xinxiang Song, Yanbing Lu, Yuanchun Tang, Nini Yuan, Suhong Sun, Yingjie An, Wenqun Ding, Xing Sun, Yanrong Wei, Shuzhen Zhang, Yannong Dou, Yun Zhao, Luyao Han, Qianhua Zhu, Junfeng Xu, Shiwen Wang, Hanbo Chen, Dan Wang, Yinqi Bai, Yikai Liang, Yuan Liu, Mengni Chen, Chun Xie, Binshi Bo, Mei Li, Xinyan Zhang, Wang Ting, Zhenhua Chen, Jiao Fang, Shuting Li, Yujia Jiang, Xing Tan, Guolong Zuo, Yue Xie, Huanhuan Li, Quyuan Tao, Yan Li, Jianfeng Liu, Yuyang Liu, Mingkun Hao, Jingjing Wang, Huiying Wen, Jiabing Liu, Yizhen Yan, Hui Zhang, Yifan Sheng, Min Han, Shui Yu, Xiaoyan Liao, Xuyin Jiang, Guangling Wang, Huanlin Liu, Congcong Wang, Ning Feng, Xin Liu, Kailong Ma, Xiangjie Xu, Tianyue Han, Huateng Cao, Huiwen Zheng, Yadong Chen, Haorong Lu, Zixian Yu, Jinsong Zhang, Bo Wang, Zhifeng Wang, Qing Xie, Shanshan Pan, Chuanyu Liu, Chan Xu, Luman Cui, Yuxiang Li, Shiping Liu, Sha Liao, Ao Chen, Qing-Feng Wu, Jian Wang, Zhiyong Liu, Yidi Sun, Jan Mulder, Huanming Yang, Xiaofei Wang, Chao Li, Jianhua Yao, Xun Xu, Longqi Liu, Zhiming Shen, Wu Wei, Yan-Gang Sun

## Abstract

A comprehensive atlas of genes, cell types, and their spatial distribution across a whole mammalian brain is fundamental for understanding function of the brain. Here, using snRNA-seq and Stereo-seq techniques, we generated a mouse brain atlas with spatial information for 308 cell clusters with single-cell resolution involving over 6 million cells as well as for 29,655 genes. We have identified new astrocyte clusters, and demonstrated that distinct cell clusters exhibit preference for cortical subregions. In addition, we identified 155 genes exhibiting regional specificity in the brainstem, and 513 long non-coding RNA exhibited regional specificity in the adult brain. Parcellation of brain regions based on spatial transcriptomic information showed large overlap with that by traditional method. Furthermore, we have uncovered 411 transcription factor regulons with spatiotemporal specificity during development. Thus, our study has discovered genes and regulon with spatiotemporal specificity, and provided a high-resolution spatial transcriptomic atlas of the mouse brain.

## Introduction

The high complexity of the mammalian brain, characterized by diversity of cell types and their specific distribution and connectivity, poses a great challenge to comprehensively understand the circuit basis underlying various animal behaviors. Enormous efforts have been made to construct informative brain atlas for mice, the most widely used animal model, with great progress having been made. Anatomical brain atlas was generated by traditional cytoarchitectural features in the mouse brain^1^. In addition, a mouse brain atlas of the expression of ∼20,000 genes was also constructed with *in situ* hybridization (ISH)^2^. Moreover, a cell atlas of mouse brain produced by single-cell transcriptome sequencing has discovered the diversity of brain cell types^3^. However, a comprehensive brain atlas across the whole mouse brain with single cell resolution and high spatial resolution is still lacking.

Recently, the development of spatial transcriptomic technologies has provided an unprecedented opportunity for mapping the distribution of gene expression as well as cell types with high spatial resolution and level of completeness. Several groups have independently established spatial molecular and cellular maps of mouse brain with *in situ* hybridization-based, *in situ* sequencing-based, or *in situ* spatial barcoding-based technique^4^. Multiplexed error-robust fluorescence *in situ* hybridization (MERFISH)^5^ and spatially-resolved transcript amplicon readout mapping (STARmap)^6^ exhibited high efficiency in capturing gene transcripts. However, these methods can only detect a set of pre-selected genes with limited numbers. In contrast, *in situ* spatial barcoding-based techniques such as Visium^7^ and Slide-seq^8,9^ could reveal spatial distribution of whole-transcriptome but without single-cell resolution. Currently, although many spatially resolved mouse brain atlases have been constructed by applying either image-^10–13^ or sequencing-based methods^14–16^, a comprehensive spatial map of the mouse brain with genome-wide coverage and cellular resolution was still not available.

Spatiotemporal profiling to determine genetic programs and induction of gene expression that drive brain development not only requires time series but also a high degree of completeness to enable temporal reconstruction of proliferation and differentiation of cell types. Development of the brain is guided by intrinsic genetic programs governed by a variety of specific transcription factors (TF). Previous studies have revealed molecular heterogeneity of the mouse brain at the embryonic stage with single-cell RNA and spatial transcriptomic sequencing^17,18^. However, genes with gradient distribution and time-dependent dynamics during development, potentially involved in development, were not systematically examined. Additionally, the maturation process of the brain sustained after birth, but spatiotemporal profile of gene expression during these stages were not well examined. Therefore, further analysis of spatiotemporal dynamics of genes from embryonic to adult stage is essential to comprehensive understanding the mechanism underlying brain development.

Accumulating evidences have shown that non-coding RNAs (ncRNAs), especially long non-coding RNAs (lncRNAs), play important roles in a wide range of biological processes of the mammalian brain, including brain development, maturation, and disease^19,20^. For example, lncRNA *Pnky* has been shown to be essential for brain development^21^. lncRNA *Evf2* can function as a *Dlx2* transcriptional coactivator and enhance the activity of *Dlx2*, which is essential for the differentiation and migration of neurons in the brain^22^. For some lncRNA, their functional role is highly related to their region-specific expression pattern in the brain^23^. Although bulk sequencing studies have characterized the lncRNAs in different brain regions of the mouse brain^24,25^, a systematic analysis of region-specific expression patterns of lncRNAs and regulation of the lncRNAs in regulon and their adjacent pairs throughout the brain has been lacking.

Here, we applied Stereo-seq, a spatial transcriptomic technology with single cell resolution to hundreds of brain sections^18^, and combined with single-nucleus RNA sequencing (snRNA-seq) data to generate a 3D single-cell transcriptomic atlas that illustrated the distribution of cell types across the whole mouse brain. Moreover, with the data of mouse brains at different developmental stages, we also studied the spatiotemporal dynamics of genes, gene modules, and TF regulons. This comprehensive dataset thereby provides a valuable resource of spatial atlas of the mouse brain, laying the foundation for the study of development, function, and gene regulation of the mouse brain.

## Results

### Spatial transcriptomic analysis of the mouse brain

We employed Stereo-seq, a sequencing-based genome-wide and high-resolution spatial transcriptomic technology that utilizes DNA nanoballs (DNBs) with 500 nm distance (center to center)^18^, to generate a spatial gene and cellular atlas of the mouse brain. We prepared coronal sections (10-µm thick) of the left hemisphere of the adult mouse brain at 100-µm intervals (Figure 1A). We obtained a dataset of 123 sections after sequencing and quality control, with gene expression profiles across 29,655 genes, covering 95.5% of annotated protein coding and non-coding genes. The obtained distribution maps of region and cell type specific genes by Stereo-seq were in line with ISH-based data presented in the Allen Brain Atlas (Figure S1A). To achieve single-cell resolution for Stereo-seq data, we defined the boundary of each cell based on the image of single strand DNA (ssDNA)^18^. A total of 4,229,623 cells were characterized in one mouse brain, with averaged molecular identifiers (MIDs) of 1,267 and averaged detected genes of 668 for each cell (Figure S1B, Table S1). Moreover, we manually segmented the brain areas for each section based on cytoarchitectural pattern and brain region delineation in Allen Mouse Brain (ABA) Common Coordinate Framework (CCFv3) (Figure 1A).

**Figure 1.**
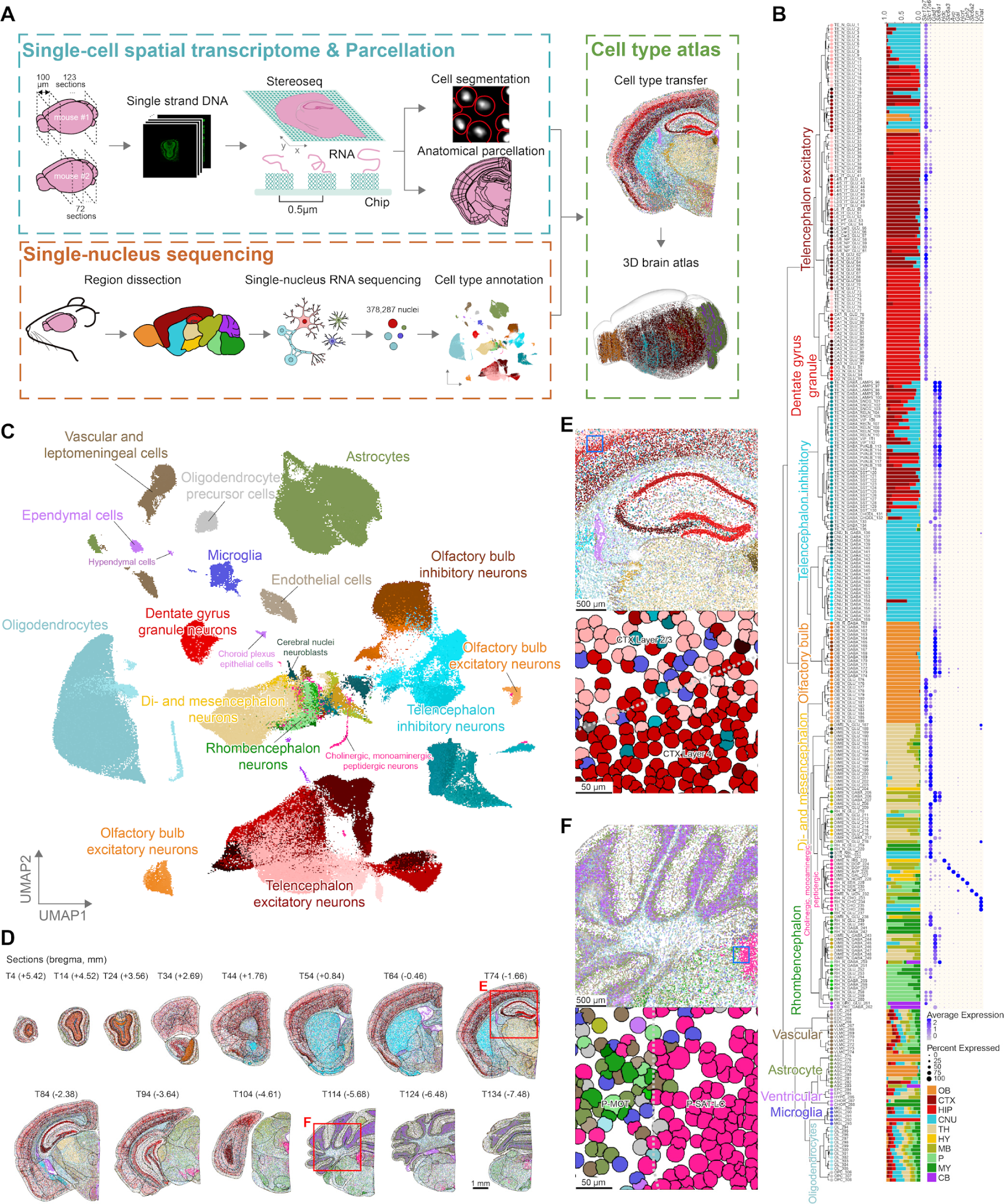
Construction of cell-type atlas across mouse brains in high resolution. **(A)**Schematic procedures showing construction of the spatial transcriptome for the whole mouse brain. Two mouse brains were sectioned every 100 µm. The red circles indicate cell boundaries. Anatomical regions were annotated according to Allen CCFv3 parcellations. **(B)** A taxonomy tree of 308 cell clusters of the mouse brain based on snRNA-seq data. Each cluster was assigned to a unique cluster name and color, attached to the end of each branch. A bar plot represents the fraction of cells according to region sources of snRNA-seq clusters (OB, olfactory bulb; CTX, cerebral cortex; HIP, hippocampus; CNU, cerebral nuclei; TH, thalamus; HY, hypothalamus; MB, midbrain; P, pons; MY, medulla; CB, cerebellum). Abbreviations of mouse brain areas were listed in Table S10. Dotplot shows the expression of neurotransmitter/neuromodulator-related genes for neurons in snRNA-seq data (*Slc17a7* and *Slc17a6*, glutamate; *Gad1* and *Slc6a1*, GABA; Hdc, histamine; *Slc6a3*, dopamine; *Avp*, arginine vasopressin; *Gal*, galanin; *Hcrt*, hypocretin; *Tph2*, Serotonin; *Slc6a2*, norepinephrine; *Ucn*, urocortin; *Chat*, Acetylcholine). **(C)** Cell subclasses and clusters in the UMAP space. **(D-F)** Overview of cell-cluster distribution of representative coronal sections in mouse brain. **(D)** Fourteen representative sections were selected every 1 mm in the brain of mouse #1 (Section No. and bregma coordinates shown, unit mm). Scale bar, 1 mm. **(E-F)** Zoomed-in views of two regions labeled by red boxes in **(D)**. Scale bar, 500 µm. Boundaries of cortical layer 4 **(E)** and locus ceruleus **(F)** are labeled by blue boxes and further magnified to visualize individual cells clearly. Scale bar, 50 µm.

Given that the average number of detected genes for each cell was not enough for precise cell type annotation, we thus performed snRNA-seq for all major brain regions of whole mouse brains, including the olfactory bulb, cerebral cortex, hippocampus, cerebral nuclei, thalamus, hypothalamus, midbrain, pons, medulla, and cerebellum (Figure 1A), in order to obtain a comprehensive and precise set of cell types. We collected 378,287 high quality nuclei and classified them into 6 cell classes, 18 cell subclasses and 308 cell clusters by iterative clustering (Figure 1B, 1C, Table S2). We further manually annotated cell clusters using canonical marker genes^3,5,26^ (Figure S1C), and obtained 262 neuronal clusters (9 subclasses), 9 astrocyte clusters (1 subclass), 5 microglia clusters (1 subclass), 15 oligodendrocyte clusters (2 subclasses), 5 ventricular clusters (3 subclasses), and 12 vascular clusters (2 subclasses). To evaluate the robustness of these cell clusters, we used a random forest model to test the classification accuracy on all clusters. The confusion matrix of the model showed high average classification accuracy (83.1%), indicating high robustness of cell clusters (Figure S1D). In addition, we compared our cell clusters with two previously published mouse brain datasets^3,26^, and found that the cell clusters in our results have a high degree of overlap with these published results (Figure S1E), indicating a high degree of consistency.

We next annotated cells sequenced with Stereo-seq by leveraging the cell clusters defined in the snRNA-seq dataset. This was achieved by utilizing Spatial-ID, a graph convolution network (GCN)-based method^27^, to assign cell clusters to each individual cell on the coronal sections. The transfer of all 308 cell clusters was conducted to over 4.2 million high-quality cells, defined as cells with more than 100 genes expressed in the Stereo-seq. The distribution patterns of these cell clusters were demonstrated in 14 selected coronal sections (spacing 1 mm apart), illustrating the continuity of various cell types along the anterior to posterior axis of the mouse brain (Figure 1D). Some cell clusters exhibited distribution to restricted brain areas. For example, we observed laminar-distributed telencephalon excitatory neurons (colored in red) in the hippocampal formation and cortex (Figure 1E), and area-specific noradrenergic neurons (colored in pink) in locus coeruleus (Figure 1F). Next, we tested technical and biological reproducibility of our results by collecting 72 coronal sections from another mouse brain covering the midbrain to hindbrain (mouse #2, Figure 1A, S1B). Comparing sections with similar bregma coordinates from two animals, revealed a consistent distribution of cell types and molecular signatures (Figure S1F-H).

By integrating our 123 coronal sections dataset with ABA CCFv3, we constructed a comprehensive 3D atlas of cell type distribution in the mouse brain (Figure 1A, Movie S1). To facilitate easy access and exploration, we created an interactive website that enables users to search for the spatial distribution of specific genes or cell types within the mouse brain (https://mouse.digital-brain.cn/spatial-omics, Figure S2).

### Spatial distribution of diverse cell types in the brain

To quantitatively analyze the whole brain distribution of all 308 cell clusters, we calculated the relative distribution of each cell cluster in 66 annotated brain segments covering the entire mouse brain. As shown in the brain-wide distribution of these cell clusters (Figure 2A), most cell clusters, especially neuronal clusters, exhibited clear regional enrichment. This is exemplified by the distribution of 10 example cell clusters, each with a distinct distribution pattern (Figure 2B, C, Figure S3A, Movie S1). For example, DG_N_GLU_92, a cluster of dentate gyrus granule cells, exhibited preferential distribution in the hippocampal region, while CNU_N_GABA_154, a cluster of medium spiny neurons, was selectively enriched in striatum.

**Figure 2.**
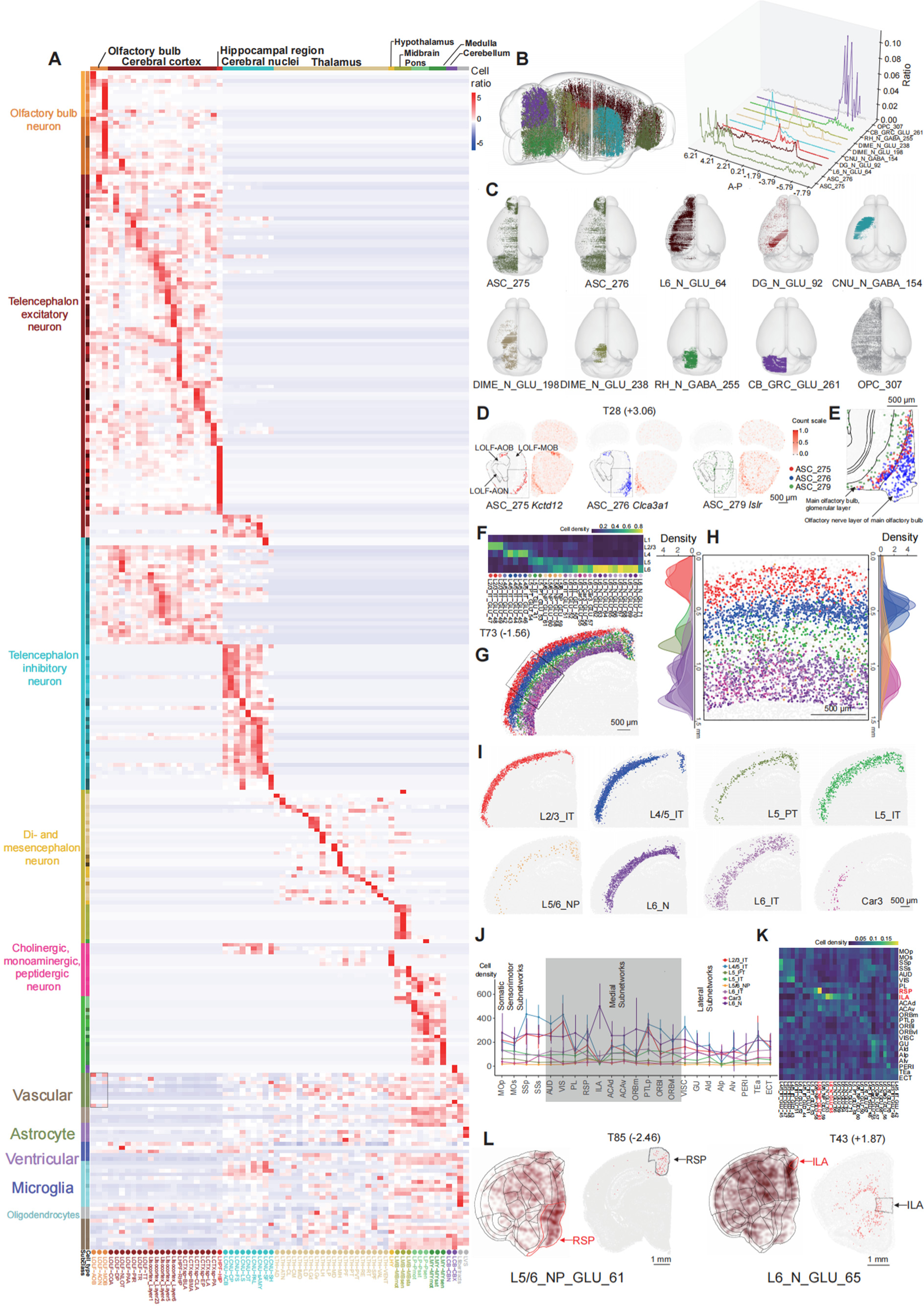
Brain-wide distribution of cells in the mouse brain. **(A)**A heatmap showing the distribution of cell clusters (308 cell clusters) across 66 brain areas. Heatmap colors the scaled cell ratio of each cluster across brain areas. **(B)** Distribution of 10 example cell clusters along anterior to posterior axis. 3D distribution of 10 representative cell clusters in the whole mouse brain (left panel). Each color represents one cell cluster. Ratio distribution of 10 example cell clusters from anterior to posterior (right panel). **(C)** Top view of 10 representative cell clusters across the whole mouse brain. **(D-E)** Distributions of cell clusters and expression of marker genes of 3 representative astrocytes cell clusters (ASC_275, ASC_276, ASC_279) individually. **(D)** ASC_275 are in red, ASC_276 are in blue, ASC_279 are in green, and other cells are in gray. Scale bar, 500 µm. **(E)** Zoomed in view of gray rectangle regions of panel D. Scale bar, 500 µm. **(F)** Heatmap of the scaled density distribution of cell clusters from 8 representative excitatory cell groups across layers of the isocortex. **(G-I)** The distribution of excitatory neurons in a coronal section of T73. **(G)** Each color represents one cell cluster. Scale bar, 500 µm. **(H)** Zoomed in view of the black rectangle region in panel G. The cortical depth distributions of individual excitatory cell clusters are also shown on the left and right sides. Scale bar, 500 µm. **(I)** The distribution of 8 excitatory cell groups as panel F in the coronal section of T73 individually. Scale bar, 500 µm. **(J)** Heatmap of the scaled density distribution of cortical excitatory neuron cell clusters across cortical areas. **(K)** Cell density of the 8 excitatory cell groups across anatomical regions of the isocortex arranged in 3 subnetworks based on their anatomical connectivity^86^, error bars were estimated across all cortical sections (mean ± SD). **(L)** Cell clusters distribution of L5/6_NP_GLU_61 and L6_N_GLU_65 on flatmap (left) and the coronal section of T85 and T43 (right). RSP and ILA regions are highlighted. Scale bar, 1 mm.

Although the non-neuron cells were widely distributed compared to neurons, some non-neuronal cell clusters exhibited specific distribution patterns (Figure 1B, 2A, Figure S3B-C, Table S3). In fact, among 9 cell clusters of astrocytes, ASC_281 (*Agt*+) specifically distributed in non-telencephalon areas; ASC_283, identified as a Bergmann glial^28^ cell due to its *Gdf10* expression profile, exhibits a distinctive localization within the cerebellum^3^ (Figure S3D-F). In addition, we found that 3 clusters (ASC_275, ASC_276, and ASC_279) of astrocytes exhibited distinct distribution patterns in subregions of olfactory bulb (Figure 2D, Figure S3C). ASC_275 and ASC_276 were predominantly localized in the regions of the accessory olfactory bulb (AOB) and the main olfactory bulb (MOB), respectively. In contrast, ASC_279 primarily resided within the MOB region. In the MOB and its surrounding area, ASC_279 predominantly resided in the glomerular layer, whereas ASC_276 was primarily situated in the olfactory nerve layer. ASC_275, on the other hand, was positioned at the boundary between the two layers (Figure 2E, Figure S3C). These 3 clusters of astrocytes showed distinct molecular signatures: ASC_275 highly expressed *Kctd12*, ASC_276 highly expressed *Clca3a1* and ASC_279 highly expressed *Islr* (Figure S3E).

The extensive coverage of our data including the cerebral cortex provides an unprecedented opportunity for examining layer and regional distribution of cells from relevant clusters. Most excitatory neurons are organized regional and layer-specific distribution patterns (Figure S3G). Among the excitatory neuronal clusters defined in snRNA-seq data of this study, 31 clusters were highly overlapped with the published single cell dataset of the mouse cerebral cortex^26^ (see Methods). These selected clusters were further categorized into 8 cell groups according to their transcriptome similarity with previously defined cortical layer specific cell types (Figure S3H). The distribution patterns of the 31 excitatory cell clusters in cortex showed laminar distribution in Stereo-seq data, consistent with a previous study^5^ (Figure 2F-I, Figure S3I). In addition, the positioning of these excitatory neurons was in agreement with the expression patterns of their marker genes, which is also consistent with the ISH data from Allen Brain Atlas (Figure S3J). In addition, the distribution pattern of different excitatory neuronal clusters across various cortical areas revealed several cortical area-specific neuronal clusters (Figure 2J). A set of specific types of neurons on layer 6 was significantly enriched in the infralimbic area (ILA) compared to other cortical areas. Quantitative analysis identified 4 cell clusters (L6_N_GLU_63, L6_N_GLU_65, L6_N_GLU_66 and L6_N_GLU_69) that are specifically enriched in the layer 6 of ILA (*p* values < 0.001) (Figure 2K-L and Figure S3I, K). Moreover, L5/6_NP_GLU_61 mainly populated the retrosplenial area (RSP) (Figure 2K-L).

Besides, we analyzed the distribution patterns of inhibitory neuron clusters in the cortex. From the classification of snRNA-seq data, 34 GABAergic inhibitory neuronal clusters in the cortex were divided into seven groups according to previous studies showing that the expression of *Lamp5*, *Sncg*, *Reln*, *Vip*, *Pvalb*, *Sst*, and *Chodl* genes, marking key GABAergic neuron subtypes^26,29^ (Figure S3L). The inhibitory neuronal clusters on Stereo-seq data showed that most inhibitory neurons in the cortex exhibited less cortical layer preference (Figure S3M), while several clusters still had layer specificity. For example, Lamp5 inhibitory neurons (including 5 cell clusters) were enriched in the superficial layers and Sst inhibitory neurons (including 11 cell clusters) were enriched in the deeper layers. We further investigated whether inhibitory neurons exhibited regional preference for distinct cortical areas. By calculating the cell densities of GABAergic inhibitory neuronal clusters in different cortical areas, several cell clusters with specific cortical region preference were identified (Figure S3N-S). For example, Pvalb neurons were rare in the agranular insular area (AIP) while Sst and Reln neurons were enriched in the ILA area (Figure S3O). Compared with Sst neurons, Pvalb neurons had significantly lower cell densities in Lateral Subnetworks and some Medial Subnetworks areas, whereas they had higher densities in the RSP (Figure S3P-R). Among all Pvalb clusters, 3 Pvalb clusters (TE_N_GABA_PVALB_115, TE_N_GABA_PVALB_116, TE_N_GABA_PVALB_118) showed this pattern. In addition, we observed that Pvalb and Sst neurons were dominant in the mouse cortex compared to other GABAergic neurons (Figure S3T-U).

To summarize, our data presented fine-grained cell type landscapes of the mouse brain. This atlas demonstrated that neuron cells are more regional-specific compared to non-neural cells, including the well-known laminar distributions of excitatory neurons and also the regional specific inhibitory neurons in cortex. Additionally, our atlas illustrated that the spatial distribution of different types of astrocytes varies across brain regions. Furthermore, we observed subtle differences in the distribution locations of three astrocyte clusters (ASC_275, ASC_276, ASC_279) in the glomerular or nerve layer of the olfactory bulb, indicating the functional variations of astrocytes.

### Distribution of the neuronal subtypes and region-specific genes in the brainstem

The brainstem is critical for multiple physiological functions, including breathing, awareness, movement^30^. However, the cell type distribution in the brainstem is not clear. In our snRNA-seq dataset, we identified 23 neuronal cell clusters in the brainstem. By combination with stereo-seq data, we found that several cell types exhibited regional specificity in the brainstem (Figure 3A). We confirmed that spatial distribution of nuclei-specific cell types was consistent between two mice examined. For example, RH_N_GABA_257, RH_N_GLU_210, RH_N_GLU_240, and RH_N_GLU_252 were restricted in the nucleus of the trapezoid body (NTB), the parabrachial nucleus (PB), the principal sensory nucleus of the trigeminal (PSV), and the cochlear nuclei (CN), respectively (Figure S4A). Furthermore, both NTB and CN are auditory nuclei, and it was intriguing that the corresponding cell types GABA_257 and GLU_252 neurons in these two nuclei were both positive for parvalbumin (coding by *Pvalb* gene) (Figure S4B). In contrast, non-neuronal cell clusters didn’t show obvious region-specific distribution in the brainstem (Figure 2A).

**Figure 3.**
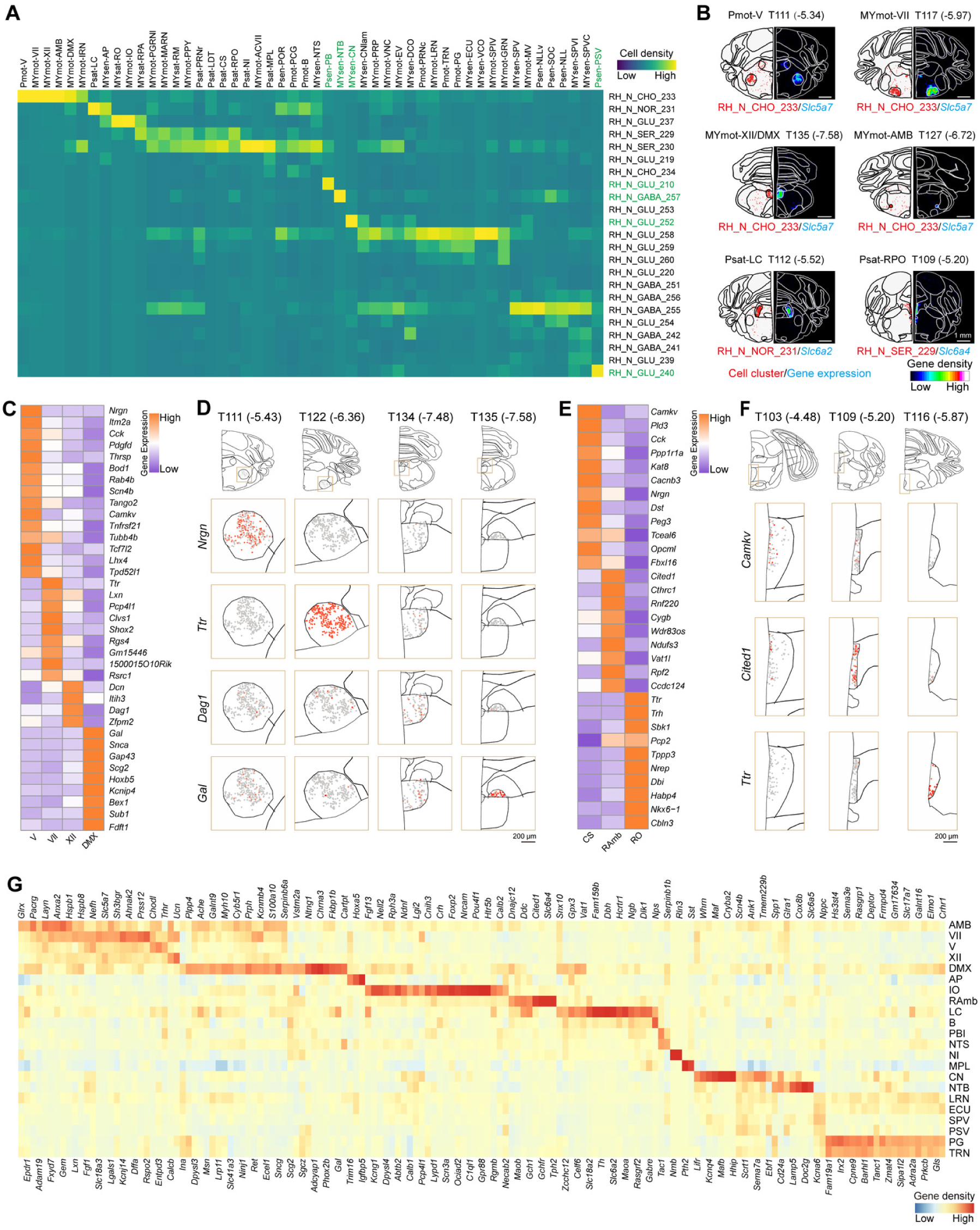
Region-selective neuronal cell clusters and genes in the brainstem. **(A)**Heatmap showing the density of different cell types in the brainstem. Horizontal and vertical axes indicate brainstem nuclei and cell clusters, respectively. **(B)** Spatial distribution of region-selective cell clusters (red) and marker genes (blue). Red dots in the left hemisphere section indicate distribution of cell clusters. The number in the parentheses on the top of each brain section represents the distance from bregma point. Scale bar, 1 mm. **(C)** Heatmap showing the expression of genes selective for 4 motor nuclei. **(D)** Spatial visualization of 4 representative genes of the 4 motor nuclei. The number in the parentheses on the top of each brain section represents the distance from bregma point. Scale bar, 200 µm. **(E)** Heatmap showing the selective expression of genes in 3 raphe nuclei. **(F)** Spatial visualization of 3 representative genes of the 3 raphe nuclei. The number in the parentheses on the top of each brain section represents the distance from bregma point. Scale bar, 200 µm. **(G)** Heatmap showing the 155 region-specific genes in the brainstem.

Motor neurons and neuromodulatory neurons are enriched in the brainstem. Consistently, our analysis also detected cholinergic (RH_N_CHO_233), norepinephrinergic (RH_N_CHO_231) and serotonergic (RH_N_CHO_229) neurons that were enriched in the motor nuclei, the locus coeruleus (LC), and the raphe nuclei, respectively (Figure 3A, 3B, Movie S2). There are multiple motor nuclei in the brainstem. It is interesting to know whether these motor nuclei have distinct molecular signatures. This has not been possible due to their small size, diffuse boundaries of these nuclei and consequently lack of whole-transcriptome sequencing data without contamination from surrounding area. We took advantage of the spatial transcriptomic data, and analyzed transcriptomic property of the motor nucleus of trigeminal (V), the facial motor nucleus (VII), the hypoglossal nucleus (XII), and the dorsal motor nucleus of the vagus nerve (DMX). We found that the four motor nuclei exhibited high heterogeneity (Figure 3C, Figure S4C). Many genes exhibited preference among 4 motor nuclei. For example, *Nrgn*, *Ttr*, and *Gal* were predominantly expressed in V, VII, and DMX, respectively, and *Dag1* were enriched in XII (Figure 3D). Gene ontology (GO) analysis also revealed differences among the four motor nuclei (Figure S4D). In addition, we performed similar analysis for serotonergic neurons in distinct raphe nuclei, and found that different raphe nuclei exhibited selective expression for specific genes (Figure 3E, F). Thus, the spatial transcriptomic analysis revealed gene expression diversity for cholinergic and serotonergic neurons with distinct spatial location.

The high resolution spatial transcriptomic data provided an opportunity for identification of regional specific genes (see Methods). We have identified 155 region-selective genes, and these genes were located in 22 brainstem nuclei (Figure 3G, see Figure S4E for examples). For example, *Pth2* and *Sst* were enriched Medial paralemniscal nucleus (MPL), consistent with previous observation^31^. Further, we have classified these regional specific genes, and found the top-ranking categories of those regional-specific genes coding enzymes, receptor-related proteins, neuropeptides, and transporters (Figure S4F, Table S4). Notably, *Dbh*, encoding the oxidoreductase which catalyzes the synthesis of norepinephrine, was selectively expressed in the LC, *Ddc* and *Tph2* that involved in serotonin biosynthesis were enriched in the Ramb, while multiple enzyme related genes such as *Plpp4*, *Dpysl3*, *Ache*, *Galnt9*, *Cyb5r1*, *Ecel1*, and *Fkbp1b* were enriched in the DMX. Several neuropeptides (*Nmb*, *Nps*, *Pth2*, *Rln3*, *Ucn*) exhibited high region-specificity in the brainstem (Movie S3). In addition, we found that some of these genes exhibited subregional specificity. For example, *Barhl1* and *Hhip* were enriched in the dorsal and ventral cochlear nucleus (CN), respectively (Figure S4G). Together, our analysis provided a map for the 23 neuronal cell clusters and revealed many region or subregion-specific genes in the brainstem.

### Gene-based brain region parcellation and spatial distribution of gene modules

Traditional segmentation of brain areas was based on cytoarchitecture and function, while the transcriptional properties of different brain regions have not been taken into account. With the brain-wide spatial transcriptomic map of the mouse brain, we examined whether brain region parcellation could be achieved based on spatial transcriptomic map. To this end, we generated a brain atlas of molecularly defined brain regions by using bin100 data (100 × 100 DNB spots, dimension size: 50 µm × 50 µm) from Stereo-seq for clustering in brain sections of mouse #1 (divided into 12 groups, see Methods). Specifically, the bin100 spots were classified by Louvain clustering, and many bin100 clusters exhibited region specific distribution, as shown in Figure S5A. We further determined the brain region occupied by each cluster based on their molecular profile and spatial location (see Methods). We found that these bin100 clusters were mainly distributed in 57 brain areas or cortical layers illustrated with distinct colors (Figure 4A). We evaluated the consistency between brain regions defined by transcriptomic data and that of ABA CCFv3 in 14 representative brain sections, and found that 70% of them coincided well with a mean Jaccard index of 0.8 (Figure S5B). This is exemplified in section T74 showing that the cortex (CTX), hippocampus (HIP), thalamus (TH), and caudate putamen in striatum (STRcp) defined by transcriptomic data exhibited high overlap with ABA CCFv3 (Figure 4B and 4C). Similarly, there is a high degree (mean Jaccard index: 0.81) of similarity for subregions of the cortex, thalamus, hippocampus, and striatum (Figure 4D). Although most of the brain regions we defined were consistent with that in the ABA CCFv3, the parcellation based on transcriptomic data could further divide some brain regions into subregions. For example, our result showed that the granular layer of the olfactory bulb (MOBgr) could be divided into medial and lateral parts in 3D space (Figure 4E). This is further confirmed by the data showing that these two areas expressed distinct molecular markers (Figure 4F). In addition, we further examined the reproducibility of brain region segmentation across 3 similar brain sections. Two of them were adjacent sections from mouse #1, and the third one was from mouse #2 with almost the same bregma coordinate. Co-clustering results on these brain sections showed that bin100 clusters exhibited similar spatial patterns across these three brain sections (Figure S5C), and that the gene expression profiles of corresponding clusters were highly correlated (Figure S5D). These results showed that brain region parcellation could be achieved based on spatial transcriptomic features.

**Figure 4.**
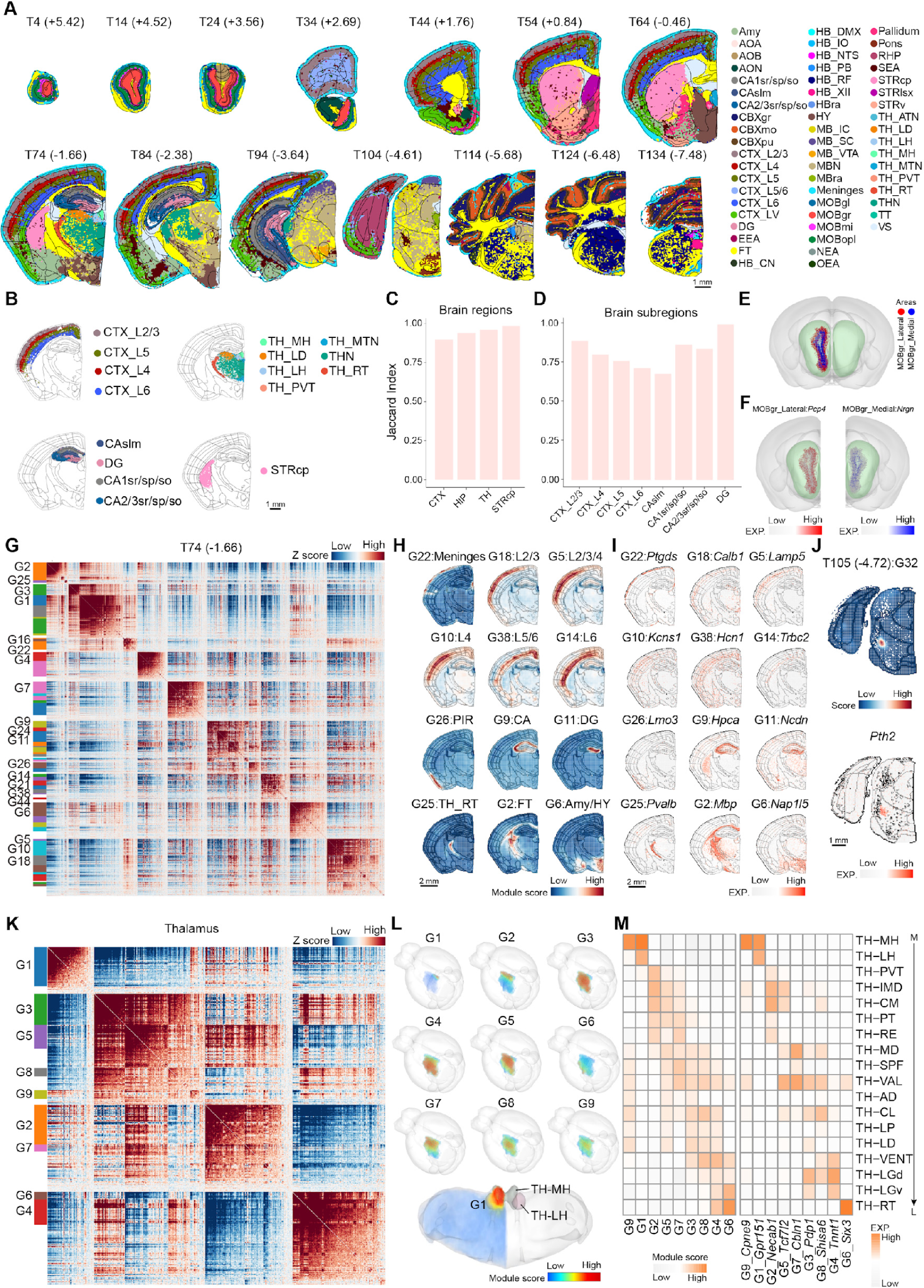
Spatial gene profiles of adult mouse brain. **(A)**Spatial visualization of brain regions determined by spatial gene profile in 14 representative brain sections along anterior to posterior axis of mouse #1 brain. Each color in brain sections represents one brain region. The black lines in each brain section are region boundaries based on ABA CCFv3. The number on the top of each section represents the distance from bregma point. Scale bar, 1 mm. **(B)** Spatial visualization of brain regions within the T74 brain section by clustering bin100 spots. Scale bar, 1 mm. **(C-D)** The histogram quantified the degree of similarity between molecular defined brain areas and corresponding brain regions delineated based on ABA CCFv3. **(C)** Jaccard similarity coefficients for CTX, TH, HIP, and STRcp brain regions. **(D)** Jaccard similarity coefficient for brain subregions in CTX and HIP. **(E)** Spatial visualization of the parcellation of the MOBgr region in 3D mouse brain. **(F)** Expression of feature genes in the granular layer of the medial and lateral olfactory bulbs. **(G)** Spatial visualization of brain region-related gene modules. Heatmap showing the genes with significant spatial autocorrelation grouped into different gene modules based on pairwise spatial correlations in T74 brain sections. The heatmap highlights 20 spatially specific gene modules. G, gene module. **(H)** Spatial visualization of 16 spatially specific gene modules. Scale bar, 2 mm. **(I)** Spatial visualization of representative genes in 16 spatially specific gene modules. Scale bar, 2 mm. **(J)** Spatial visualization of a gene module enriched in the MPL and the expression of a representative gene in this module. Scale bar, 1 mm. **(K)** Heatmap showing the genes with significant spatial autocorrelation grouped into different gene modules based on pairwise spatial correlations in the whole thalamus. **(L)** 3D visualization of 9 gene modules related to the subregion of thalamus (upper) and highlighting that module 1 is primarily associated with the medial habenula in the thalamus (bottom). **(M)** Brain region specificity of gene modules in the 3D thalamus. Left panel: module enrichment scores of the thalamic gene modules in thalamus subregions. Right panel: heatmap showing expression of representative genes of the thalamic gene modules in thalamus subregions. The subregions of the thalamus were ordered in their spatial location along medial-lateral (M-L) axis, as shown by the vertical arrow on the right.

The function of each brain area is highly dependent on the genes expressed in these brain areas. We thus examined the genes co-expressed in different brain areas across the whole brain. We analyzed the region-specific gene module which is defined as a group of genes with similar spatial patterns. We characterized gene modules in 123 sections from mouse #1 using Hotspot, a spatially-varying gene identification method^32^. We have identified 2,632 regional specific gene modules in brain sections of mouse #1, and we also summarized the brain area specific genes over the whole mouse brain (Table S5, see Methods). For example, we found 50 modules in the T74 brain section (Figure 4G), of which 34 modules exhibited regional specificity (Table S5). The distribution of 12 modules was shown in Figure 4H. Specifically, Gene module 22 (G22), G18, G5, G10, G38, and G14 were enriched in the CTX, and each with preference for different cortical layers. G26 was selectively located in the piriform cortex (PIR). Notably, G9 and G11 were specifically enriched in the ammon’s horn (CA) and the dentate gyrus (DG) regions of the HIP, respectively. G25, with *Pvalb* exhibiting the highest autocorrelation coefficient, was found in the reticular nucleus of thalamus (TH_RT), in line with aggregation of mass Pvalb GABAergic neurons in the reticular nucleus^33,34^. Brain region or subregion specific identity of different gene modules were further evidenced by specific distribution of representative genes of these gene modules (Figure 4I and S5E). The GO functional enrichment analysis showed that genes enriched in the modules were relevant to the brain region where the modules were located. For example, G2 was located in the fiber tracts, and was enriched in axon ensheathment, and G6 was located in hypothalamus, and was enriched in hormone secretion (Figure S5F). Additionally, gene module analysis also depicted small nucleus not defined in CCFv3. For example, we found a spatially-specific gene module enriched in the MPL (Figure 4J). The reliability of gene module analysis was verified by comparing the modules of adjacent brain sections, and we found high correlation between spatial gene modules in two adjacent brain sections (Figure S5G-J).

The examination of gene module distribution would likely be facilitated by exploring their distribution in a 3D manner. We thus took thalamus as an example, and analyzed all 42 sections containing the thalamus and determined their gene modules in 3D space through Hotspot^32^. We found 9 gene modules in the thalamus (Figure 4K). Some of the modules were restricted in specific thalamic regions. For example, G1 was located in the medial and lateral habenula nucleus. While some modules were distributed in several brain areas. For example, G6 was located in reticular nucleus of the thalamus (TH_RT), the ventral part of the lateral geniculate complex (TH_LGv), the dorsal part of the lateral geniculate complex (TH_LGd), and the ventral group of the dorsal thalamus (TH_VENT) (Figure 4L and 4M). We found that the module scores, the scores of bin100 spots within subregions of the thalamus for the gene sets of each gene module, and the expression patterns of representative genes within each module, reflected the spatial preferences for each module. The medial and lateral thalamus exhibited different preferences for different gene modules (Figure 4M). Thus, we have identified regional specific gene modules across the whole brain, and the examination of gene module distribution was facilitated by exploring their distribution in a 3D manner.

### Spatiotemporal profile of gene expression in the developing brain

The brain development is governed by sets of genes with distinct spatiotemporal profile. To explore the spatiotemporal dynamics of gene regulation, we collected spatial whole-transcriptome data of developing mouse brain with 7 sagittal sections from embryonic to adult stage (clustered and annotated similarly to coronal sections) (Figure 5A and S6A) including published data on E12.5, E14.5, E16.5^18^ and P7^35^ mice and newly generated data for P1, P14, and P77 mice. Given that TF regulons, where a core TF dictates a series of downstream genes, are common units for gene regulation, we thus determined spatiotemporal properties of TF regulons. By using gene regulatory network inference and clustering algorithms (SCENIC) for each of the 7 developmental time points^36^, we identified a total of 998 TF regulons for 7 sagittal sections. To examine the potential interaction between TF regulons, we determined the spatial co-localization of these TF regulons, as the TF with strong interaction should colocalize spatially. We applied Hotspot^32^ on each section, resulting in a total of 150 clusters, each of which contains a group of spatially co-localized TF regulons (Figure 5B, Table S6). GO enrichment analysis found that the regulons within each cluster were related to specific biological processes (Figure 5B), suggesting these spatial co-localized regulons could work together to empower brain-regional development or function. Interestingly, we found 144 of these clusters localized in major brain regions. For example, 6 clusters from section of P4 mouse localized in cortex (cluster 19), striatum (cluster 3), thalamus (cluster 16), cerebellum (cluster 8), olfactory bulb (cluster 6) and ventricle (cluster 10), respectively (Figure 5C).

**Figure 5.**
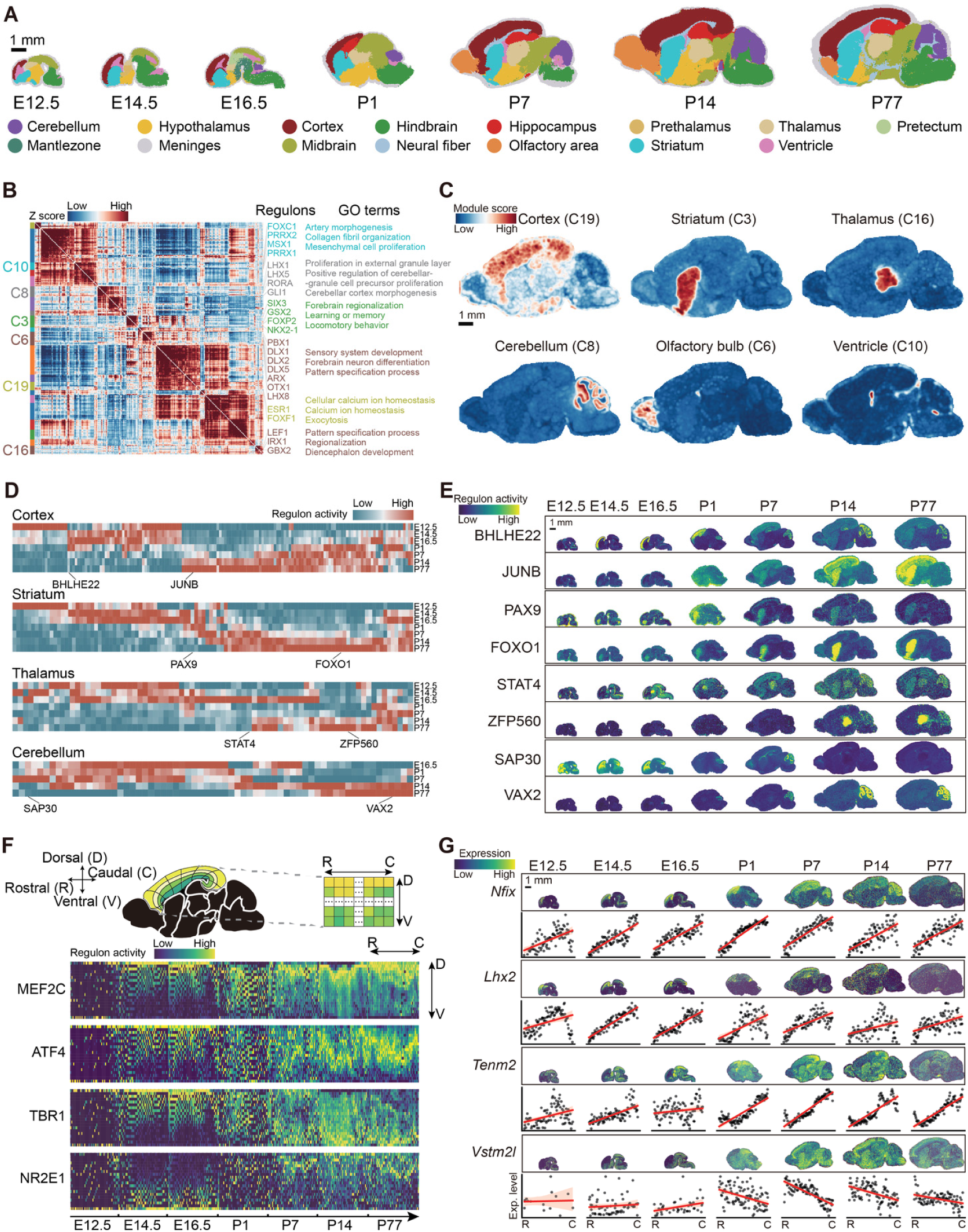
Spatial molecular profile of developmental mouse brain. **(A)**Graph showing the distribution of clusters generated by unsupervised spatial constrained clustering (SCC) of mouse brain sections across E12.5-P77. Clusters were annotated with brain-related marker genes. Scale bar, 1 mm. **(B)** Heatmap showing identified TF regulons in P14 sagittal brain section with significant spatial autocorrelation. TF regulons with high spatial correlation were grouped into clusters related to distinct gene ontology biological processes. C: TF regulon cluster. **(C)** Spatial distribution of 6 region-specific TF regulon clusters in P14 sagittal brain section. C: TF regulon cluster. Scale bar, 1 mm. **(D)** Heatmap illustrating the TF regulon activities during development in cortex, striatum, thalamus, and cerebellum. Regulon activity was calculated using AUCell in SCENIC^36^ (see Methods). **(E)** Spatial visualization of TF regulon activities of 8 TF regulons in developing mouse brain (cortex, striatum, thalamus and cerebellum) across 7 stages. Scale bar, 1 mm. **(F)** Top: Sketch of the orthogonal visualization for cortical laminar-columnar structure. Bottom: Heatmap showing the spatial gradient of 4 example TF regulons along the dorsal-ventral laminar axis in the developmental cortex (E12.5-P77). **(G)** Spatial visualization of dynamic changes of genes along the rostral-caudal gradient axis in the developing cortex. The gradient of each gene along rostral-caudal axis at each stage is demonstrated using an auxiliary scatter plot, with black dots indicating the mean expression on relative rostral-caudal position (see Methods), and a red line showing the linear regression of these points. Scale bar, 1 mm.

Next, we asked how the activities, as defined by an expression-based metric from SCENIC^36^, of these region-specific TF regulons change during development. In the cortex, 68 TF regulons exhibited higher activity during embryonic stages, while 76 TF regulons exhibited higher activity in postnatal stages (Figure 5D). In striatum, we found that 52 TF regulons were more abundant in early periods and that 56 TF regulons were abundant in later periods (Figure 5D). We also found a similar pattern for 77 and 65 TF regulons in thalamus and cerebellum, respectively (Figure 5D). For example, BHLHE22 (names in all caps stand for TF regulons) was found to be abundant in embryonic cortex and developing cerebellum (Figure 5E), where *Bhlhe22*, the core TF of this regulon, was considered to regulate the postmitotic acquisition of area identities in developing neocortex^37^. On the other hand, JUNB, the TF regulon of a well-known immediate early gene *Junb^38^*, was found abundant specifically in postnatal stages. In striatum, PAX9 was enriched in embryonic age, while FOXO1 was gradually enriched during postnatal development, consistent with previous reports in developing mouse striatum^39^ (Figure 5E). Other examples were shown in Figure 5E. Additionally, we investigated individual genes that were specifically expressed in cortex, striatum, and thalamus. Within these brain regions, 33 genes were found to be region-specific across all 7 stages, while a total of 385 genes showed brain-regional specificity only at a certain stage (Figure S6B). Collectively, a comprehensive dynamic profile of regional specific regulons and genes was summarized in Table S6. Thus, these analyses revealed TF regulons and genes showing distinct spatiotemporal properties.

Cortex was organized as both laminar and columnar structures during development, which is determined by transcriptional programs^40^. We determined transcriptional regulation across the dorsal-ventral axis or rostral-caudal axis during development using Spateo^41^. In analysis of the dorsal-ventral axis, we found 206 regulons showing laminar related distribution (Table S6). For example, ATF4 gradually shifted from the superficial layer to the middle layer (Figure 5F). TBR1 shifted from superficial layer to deep layer, in line with the prior maturation of layer 6 cortical neurons^42,43^. NR2E1 shifted from deep layer to superficial layer (Figure 5F), which aligns the important function of *Nr2e1* in neural stem cell proliferation^44,45^ with its crucial role for the intactness of supragranular layer^46^. In analysis of the rostral-caudal axis, we found 22 TFs exhibiting spatial increment or decrement gradient (Figure S6C, Table S6). For example, spatial increments along the rostral-caudal axis were found in all developmental stages for *Nfix* (Figure 5G). *Lhx2* showed a similar increment as *Nfix* in embryonic stages (E12.5, E14.5, E16.5), but the pattern was weakened in postnatal stages (P1, P7, P14, P77) (Figure 5G). This is in line with previous reports showing the gradient distribution of *Lhx2* at E15.5 stage and its crucial role in barrel column formation^47,48^, a process most active in embryonic stages^49^. In addition, we found 134 non-transcription-factor genes exhibiting rostral-caudal gradients (Figure 5G, Figure S6C, Table S6). Thus, we have revealed many TFs exhibiting gradient distribution along the dorsal-ventral axis or rostral-caudal axis during brain development.

Besides, we also examined the kinetics of general neural developmental events including gliogenesis, neuroblast proliferation, and synapse maturation using gene set enrichment scoring method (see Methods). Our analysis showed that three major neurodevelopmental event-associated gene sets exhibited the dynamic change across 9 brain regions and 7 time points (Figure S6D-F). Specifically, the genes involved in gliogenesis and synapse maturation showed higher enrichment in postnatal stages, while genes associated with neuroblast proliferation were enriched in embryonic stages (Figure S6D-E). The genes involved in gliogenesis started to show enrichment around birth and quickly reached peak level at P14. Interestingly, we found higher levels of gliogenesis-related gene enrichment in the fiber tracts and hindbrain as compared to others, consistent with the recent report in human brain gliogenesis^50^. In addition, genes involved in synapse maturation were gradually enriched at the very beginning of embryonic stages, and reached steady state after birth (Figure S6F). Thus, we have illustrated diverse kinetics of different developmental events in different brain regions.

### Spatiotemporal profile of long non-coding RNA in the brain

In the mammalian genome, less than 3% of the genome is transcribed into protein-coding transcripts, the majority is transcribed into ncRNAs, some of them with length over 200 bps, defined as lncRNAs^51^. Although previous studies have characterized the properties of a limited number of lncRNAs in the mouse brain^51,52^, there is a lack of systematic characterization of their spatial distribution pattern. Among 9,580 known lncRNAs of the mouse genome, we detected 5,834 lncRNAs in our adult mouse brain Stereo-seq dataset. Furthermore, we found that 513 lncRNAs exhibited regional specificity in distribution (see Methods, Table S7). Here, we showed the typical lncRNAs that exhibited region specificity in 14 brain regions (Figure 6A). For examples, *Gm12688*, *Gm33651*, *6330420H09Rik*, *Gm20649*, *Hotairm1* and *Gm14033* exhibited specific expression in olfactory areas (OLF), STR, HIP, hypothalamus (HY), medulla (MY), and cerebellum (CB), respectively (Figure 6B, 6C, Movie S4). Moreover, we found that 37 lncRNAs showed layer specific distribution in the cortex (Figure 6D). For example, *Gm26870*, *A830009L08Rik*, *1700047F07Rik*, *Gm11730,* and *Gm28928* was localized specifically in L1, L2/3, L4, L5, and L6, respectively (Figure 6E). Thus, we have identified the lncRNAs with distinct distribution patterns in the adult mouse brain.

**Figure 6.**
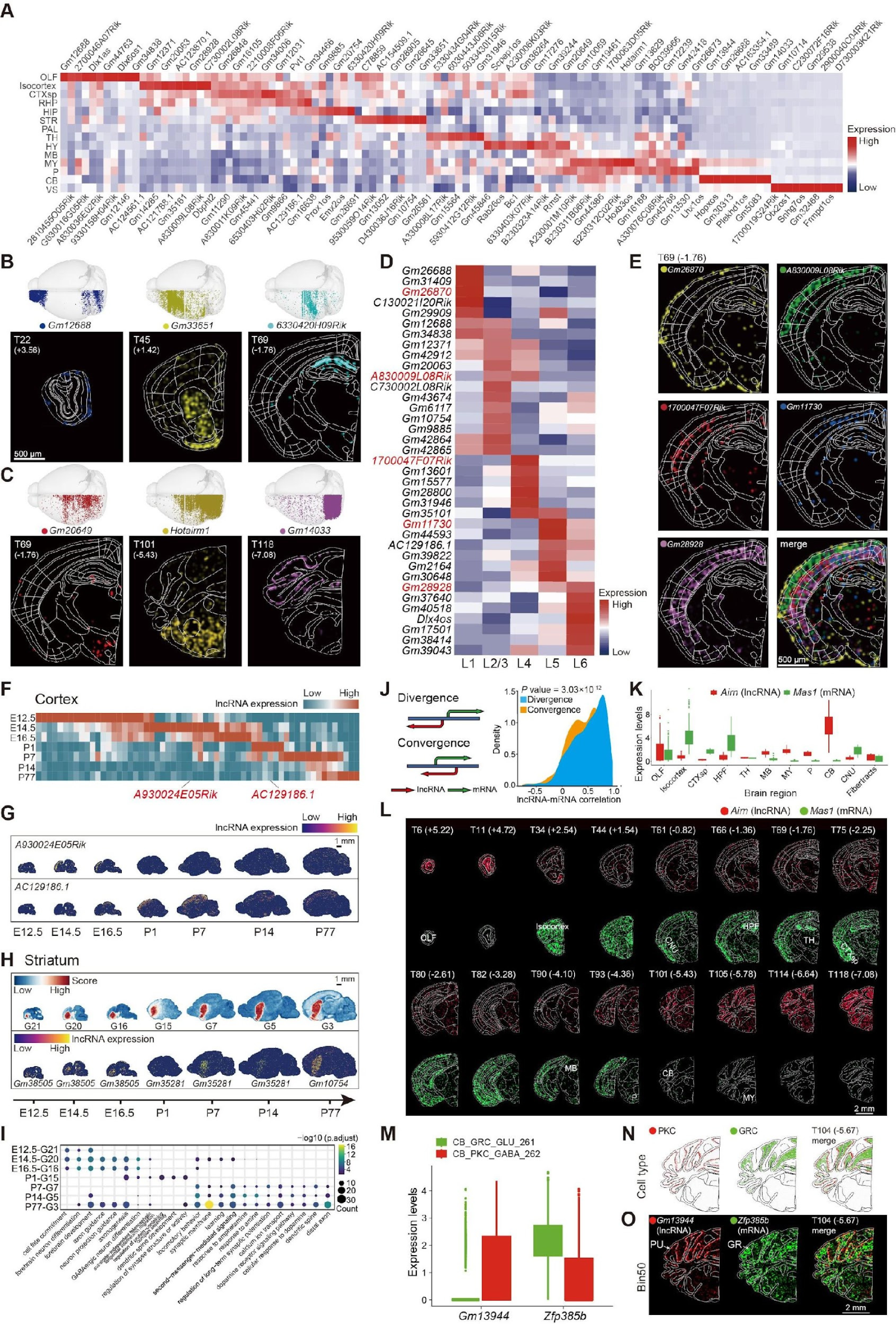
Spatial expression profile of specific lncRNA in the mouse whole brain. **(A)**Heatmap of representative region-specific lncRNAs in 14 different brain regions. **(B-C)** The whole brain distribution of expression of 6 lncRNAs (upper panel) and their spatial distribution in bin50 in related brain sections (lower panel). The number on the top of each brain section in the lower panel represents the section number and distance from the bregma point, respectively (Section No. and bregma coordinates shown, unit mm). Scale bar, 500 µm. **(D)** Expression heatmap of 37 cortical layer-specific lncRNAs. Five examples showing in panel E are colored in red. **(E)** The spatial distribution of expression in bin50 of 5 lncRNAs in the coronal section of T69. Five lncRNAs are labeled with different colors individually, and merged in the last one. Section No. and bregma coordinates shown, unit: mm. Scale bar, 500 µm. **(F)** Heatmap of the dynamic expression changes of lncRNAs in the developmental cortex. **(G)** Spatial expression distribution of spatiotemporal cortical-specific lncRNAs in 7 sagittal sections at different developmental stages (including E12.5, E14.5, E16.5, P1, P7, P14, P77). Scale bar, 1 mm. **(H)** Spatial distribution of striatum-specific modules and the expression of their corresponding lncRNAs (*Gm38505*, *Gm35281*, *Gm10754*) at different developmental stages. Scale bar, 1 mm. **(I)** Bubble plot exhibiting the representative Gene Ontology enrichment pathways of striatum-specific modules at different developmental stages. **(J)** The distribution of Pearson correlation (right) of convergent and divergent gene pairs (cartoon shown on left) in mouse brain areas. **(K)** Boxplot of the expression levels of the *Airn-Mas1* convergent gene pair in different brain regions. **(L)** The spatial expression distribution in bin50 of *Airn*-*Mas1* convergent gene pairs in 16 sections. Section No. and bregma coordinates shown, unit mm. *Airn* is colored in red and *Mas1* in green. Scale bar, 2 mm. **(M)** Boxplot of the expression levels of *Gm13944-Zfp385b* convergent gene pair in two cerebellar cell clusters of CB_GRC_GLU_261 (granule cells, GRC) and CB_PKC_GABA_262 (Purkinje cell, PKC). **(N)** The spatial distribution of *Gm13944-Zfp385b* convergent gene pairs in cell types of PKC (red) and GRC (green) in section T104. Section No. and bregma coordinates shown, unit mm. **(O)** The spatial distribution expression in bin50 of *Gm13944* (red) and *Zfp385b* (green) convergent gene pair in cerebellar Purkinje cell layer (PU) and granule cell layer (GR), respectively. White arrows mark PU and GR. Section No. and bregma coordinates shown, unit mm. Scale bar, 2 mm.

The lncRNA is known to be important for brain development^53–55^. This is mostly dependent on their regulation of coding genes. We performed Hotspot analysis of gene expression data for each section from E12.5 to P77, in order to determine the temporal dynamics of lncRNAs and coding genes with similar expression patterns. We obtained 184 gene modules, 160 of which showed brain region preference (Table S8). We showed the data for the P14 stage as an example here (Figure S7A-B). We next determined the lncRNAs with temporal specificity in the cortex, striatum, thalamus, and cerebellum across seven time points, and found that 216 lncRNAs (60 for cortex, 73 for striatum, 30 for thalamus, 95 for cerebellum) exhibited spatiotemporal specificity (Figure S7C-D). For example, in the cortex, *A930024E05Rik* was found to be highly abundant in the embryonic stage, and this gene was also shown to be specifically expressed in the developing mouse embryonic cortex^56^. While, *AC129186.1* was mainly abundant in the postnatal period (Figure 6F and 6G). Some regional specific lncRNAs revealed in our analysis have also been identified in previous studies. For example, brain region specific expression of *Gm11266* (cortex)^57^, *D430036J16Rik* (striatum)^2^, *Gm2694* (cerebellum)^58,59^, *Gm15577* (cerebellum)^60^ have been reported. For most of the lncRNAs identified in our analysis, their distribution and function were not known previously.

It has been thought that genes within the same modules are functionally relevant^61^. To predict the possible functional roles that lncRNAs may play during development, we performed GO functional enrichment analysis on brain-region specific gene modules, such as cortex, striatum, and cerebellum. We found that modules restricted at early stage (E12.5-E16.5) were related to neuronal development as evidenced by the data showing the abundance of genes related to neuronal development (Figure 6I, S7E-F). Therefore, we speculate that lncRNAs in these modules may be involved in mouse brain development (Table S8). At the late stages (P1-P77), modules begin to be enriched with genes relevant to brain region-specific functions. In the striatum, modules (including 36 lncRNAs) in the postnatal stage were enriched in the dopamine receptor signaling pathway (Figure 6H and 6I), whereas in the cortex, modules (including 15 lncRNAs) were related to maturation of functional synapses (Figure S7E). In addition, cerebellum-specific modules (including 46 lncRNAs) in the postnatal stage are associated with parallel fiber to Purkinje cell synapse among others (Figure S7F). In addition, lncRNA *Gm2694* in this module was also known to regulate synaptic stability in the cerebellum^58^. Thus, it is likely that lncRNAs may be involved in regulating brain region specific function at the late stages. These results support the notion that lncRNAs may be involved in embryonic neural development and neural function.

Recent studies have shown that lncRNA transcription occurs or is associated with nearby gene transcription, such as transcription from shared promoter regions^62–64^. Divergently transcribed lncRNAs transcribe on opposite strands from the same promoter region of their adjacent mRNAs, while convergent transcribed lncRNAs transcribe on opposite strands towards their adjacent mRNAs, often overlap each other^65^. Although the expression pattern of divergent pairs of lncRNAs and mRNAs were studied in cell lines or tissue^62,66^, the spatial expression pattern of divergent and convergent pairs in the brain was largely unknown. In order to explore the expression pattern between lncRNA and its adjacent mRNA, we divided lncRNA-mRNA gene pairs into divergent (2,503 pairs) and convergent (4,480 pairs) groups based on the distance between the transcription start sites of lncRNA and mRNA on the genome (see Methods). In our spatial transcriptomic data, we identified a total of 3,049 lncRNA-mRNA gene pairs expressed in the adult mouse brain, including 1,019 pairs of divergent and 2,030 pairs of convergent groups (Table S9). By comparing the expression of gene pairs in different brain regions, we found that the correlation of lncRNA-mRNA gene pairs in the divergent group was significantly higher than that in the convergent group (Figure 6J), with 18 of convergent gene pairs showing antagonistic expression pattern in the mouse brain (see Methods, Table S9). For example, we found that the expression levels of the convergent *Airn*-*Mas1* gene pair were negatively correlated with significant differences in diverse brain regions (Figure 6K-L, Table S9). Similar antagonistic expression pattern was also observed for another convergent gene pair *Gm45441*-*Grin2d* (Figure S7G-H). We then wanted to know whether there were differences in the spatial distribution of gene pairs among different cell types. By comparing the expression of gene pairs in different cell clusters in our spatial transcriptomic data, we found that 103 convergent gene pairs have obvious antagonistic expression in different cell types (see Methods). For example, in Purkinje cells (PKC), the expression level of *Gm13944* was higher than that of *Zfp385b*, while in granule cells (GRC), the expression level of *Gm13944* was lower than that of *Zfp385b* (Figure 6M, Figure S7I). Consistently, by examining the spatial distribution profiles in bin50 data from Stereo-seq, we also observed that specific high expression of *Gm13944* in the Purkinje cell layer (PU) but *Zfp385b* in the granule cell layer (GR) of the cerebellum (Figure 6N-O). Among these 103 gene pairs, 18 gene pairs also exhibited antagonistic expression trends in different brain regions as mentioned above (Figure S7J). Taken together, we have characterized the spatial expression pattern of lncRNA-mRNA gene pairs in the mouse brain, providing an important resource for studying the functions of lncRNAs in the brain.

## Discussion

The mammalian brain possesses a remarkably intricate organization, consisting of tens of millions to billions of cells. In this study, we have constructed a comprehensive transcriptomic and spatially-resolved cellular map of the mouse brain, and provided an atlas with genome-wide coverage with cellular resolution. We have mapped the brain-wide distribution of 308 distinct cell clusters and elucidated the region-specific of cell types and gene modules throughout the entire brain. Moreover, we identified numerous TF regulons and lncRNAs that exhibited specific spatiotemporal dynamics during brain development. These findings lay a foundation for understanding the complex interactions among diverse cell types within the brain, as well as the neural mechanism underlying animal behavior.

### Brain-wide distribution of diverse cell types

Through analysis of the spatial distribution of cell types in the mouse brain, we have discovered distinct distribution patterns for diverse cell types including excitatory neurons, inhibitory neurons, and non-neuronal cells. In the cortex, excitatory neurons exhibited a much more obvious laminar distribution compared to inhibitory neurons and non-neuronal cells, consistent with previous findings^5^. More importantly, we have demonstrated that different cortical regions exhibited preference for distinct cell types. For example, we found four cell clusters of cortical excitatory neurons were abundant in ILA, which is one of the least understood regions of the anterior cingulate cortex^67^. As ILA was reported to be specifically related in mood and affective disorder symptomatology^67^, it is likely that these ILA specific cell clusters are involved in these disorders. Similarly, we also found several clusters of inhibitory neurons exhibiting a region-specific distribution pattern such as Sst neurons in ILA. In line with our study, the abundance of Sst neurons in ILA has been reported in previous studies^68^. This early study examined the Sst neurons with transgenic labeling, and this method could only be applied to limited types of neurons. In contrast, our study identified precise cell types with transcriptomic analysis, giving us an opportunity to systematically examine their differential distribution across the whole cortex. A recent study in the cortex of macaque has demonstrated variances in cellular configurations across various cortical areas, supporting the idea that some cell types exhibit preference for distinct cortical areas^69^.

Compared with neurons, most of the non-neuronal cells are widely distributed throughout the whole brain, consistent with previous findings^3^. However, we identified several clusters of astrocytes that exhibited region-specific distribution patterns, such as ASC_283 that was predominantly localized in the cerebellum. In addition, we observed 3 clusters of astrocytes (ASC_275, ASC_276, and ASC_279) were enriched in the olfactory bulb. ASC_279 highly expresses *Slc6a11* and *Islr*, and these marker genes were found to be highly expressed in the astrocyte cell type identified in a previous study^3^, suggesting that ASC_279 is the same cell type identified previously. The other two astrocyte cell clusters showed distinct molecular signatures compared to ASC_279, indicating these two new astrocyte cell clusters. Furthermore, these three clusters exhibited distinct distribution patterns between the nerve and glomerular layers of the olfactory bulb. This distribution preference may be related to the signal processing and metabolic support to neurons on different layers of the olfactory bulb^70–72^.

In the brainstem, we observed multiple clusters of neuronal cells restricted within a certain nucleus (Figure 3A, 3B). Among these nuclei, the auditory nuclei (NTB and CN) showed high cell-type specificity. We observed that glycinergic interneurons were enriched in the NTB as documented^73^ and a specific type of excitatory neuron, RH_N_GLU_252, was predominantly confined in CN (Figure S4B). More importantly, these two clusters of neurons were both *Pvalb* positive. Pvalb interneurons are widely distributed in the auditory system, which are well-tuned for sound frequency^74^ and could enhance temporal coding in the auditory pathway^75^. The distinct distribution of excitatory Pvalb and inhibitory Pvalb neurons in different auditory nuclei, suggesting that two subtypes of Pvalb neurons play different roles in processing sound information.

### Region-specific modules and genes

Although there have been many studies on region-specific protein coding genes in the mouse brain^2,76^, those analyses were often confounded by diverse experimental sampling among different animals. In our study, all data was collected with the same approach. We have employed two approaches to identify regional specific genes. The integration of both approaches allowed us to comprehensively identify region-specific genes, especially those specific to small nuclei. We identified 155 region-specific genes in the brainstem. Some of these region-specific genes can further define fine structure of brain nuclei. For example, *Barhl1* and *Hhip* are enriched in the dorsal and ventral cochlear nuclei in the brainstem, respectively. These fine-scale organizations had not been discovered before.

Our study also identified many lncRNAs with spatial distribution specificity, with some of them having been reported in previous studies^2,58^. Furthermore, we found that some convergent gene pairs tended to be antagonistically expressed in different brain regions and different cell types in mice. We found that lncRNA *Airn* overlaps with *Mas1,* and that they have opposite expression trends in different brain regions, suggesting that *Airn* may have a potential function in silencing *Mas1* in the mouse brain. Consistently, previous studies have shown that *Airn* indeed can silence the nearby gene in mouse embryonic stem cells^77^. Notably, in the cerebellum, we found that the expression of *Zfp385b,* a zinc finger binding protein, and lncRNA *Gm13944* exhibited a complementary manner, indicating the negative regulatory effect on *Zfp385b* by *Gm13944*. This is supported by early study showing that *Zfp385b* could be regulated by lncRNAs^78^. Thus, these region specific lncRNAs could have regulatory effects on their adjacent protein coding genes in the brain.

Spatial transcriptomic dataset provides an opportunity to further determine the molecular differences among the same type of neuron distributed in distinct nuclei with small size, which is not possible with single-cell sequencing. For example, cholinergic neurons are distributed in several motor nuclei in the brain stem, and these nuclei are all small in size. By analyzing the spatial transcriptomic data, we identified nuclei-specific genes for the cholinergic neurons in discrete motor nuclei. Our GO analysis indicates that hypoglossal nuclei could be involved in several distinct physiological processes, including regulation of heart contraction, biomineral tissue development, female gonad development, lung development, and positive regulation of production of molecular mediators of immune response. This is in line with previous studies showing that perinatal lung inflammation increases proinflammatory cytokines expression in the hypoglossal nucleus^79^ and that sudden unexplained perinatal death was highly related with developmental anomalies of the hypoglossal nucleus^80^, suggesting that there may be a potential link or causal relationship between these biological processes. The identification of region-specific genes aids our understanding of the function of nuclei.

### Spatiotemporal analysis transcriptome of developing mouse brain

By leveraging the Stereo-seq technology, we constructed a whole-transcriptome map with 29,655 genes for developing mouse brain from embryonic to adult stages, extending the existing profile of around 2,100 selected genes using ISH^81^. Further analysis has identified 998 TF regulons exhibiting spatiotemporal specificity. Systemic analysis of TF regulons across different development stages has never been reported previously. Nevertheless, some of our results are supported by previous studies using either immunohistochemical or ISH methods. For example, BHLHE22 is enriched in cortex during early development, which is in line with previous studies showing that *Bhlhe22* is highly expressed in embryonic cortex and that it is involved in the identity acquisition of postmitotic cortical neurons^37^. Thus, the TF regulons showing spatiotemporal specificity are likely to be involved in brain development, although the functional roles of most regulons remain to be determined. In addition, we also discovered 385 genes showing spatiotemporal specificity. The discovery of TF regulons and genes exhibiting spatiotemporal specificity provides a valuable dataset for the study of neurodevelopment.

Further analysis showed that some of the TF regulons exhibited delicate spatiotemporal patterns within a single brain region. In the developing cortex, we showed that 206 of the identified TF regulons exhibited laminar enrichment on the dorsal-ventral axis and 156 genes showed gradient distribution on the rostral-caudal axis. Some of these findings are supported by previous findings. Our results showed that *Lhx2* exhibited gradient distribution on the rostral-caudal axis, which is consistent with a previous study^82^. This gradient distribution pattern has been thought to be important for formation of sub-structures^40^. It has been shown that *Lhx2* is involved in barrel cortex formation^47^. Thus, some of the TF regulons or genes with laminar enrichment or gradient distribution in cortex are likely to be crucial to shape cortical structure. Our study significantly extended previous findings of dozens of genes exhibiting rostral-caudal axis gradient patterns in certain developmental stages^40^, and revealed hundreds of genes exhibiting such distribution patterns across developmental stages. The function of these genes or TF regulons would need to be examined experimentally.

In the analysis of ncRNA, we observed spatiotemporal kinetics for lncRNAs in developing mouse brains. We have discovered 216 lncRNAs (60 for cortex, 73 for thalamus, 30 for striatum, 95 for cerebellum) showing specific spatiotemporal kinetics. For example, we found that *A930024E05Rik* was expressed in the cortex during early developmental stages but not later ones. Consistently, an early functional study showed that this gene is essential in proper differentiation and migration of cortical projection neurons^56^. Early studies have shown that lncRNAs could function at the transcriptional regulation and play important roles in the brain development^20^. Thus, the newly identified lncRNAs exhibiting stage-dependent expression in the brain could be involved in brain development. In line with this, our further analysis showed that some of the lncRNAs are present in the module enriched with genes relevant to neurodevelopment. This analysis of spatiotemporal kinetics in lncRNAs from embryonic stage to adult stage laid the foundation for further analysis of the function of lncRNAs.

### Parcellation of brain areas based on gene expression pattern

Traditionally, the boundaries of brain regions were defined based on cytoarchitecture and function^1,83,84^. In our analysis, we were able to define the boundaries of 57 brain areas in the mouse brain based on the spatial transcriptomic properties. The majority of brain areas we defined are consistent with the brain regions in Allen’s CCFv3^85^. Thus, the parcellation of different brain areas has their transcriptomic basis, consistent with a recent study^12^. Given that our transcriptomic data have high spatial resolution, our analysis allowed for a more detailed division of certain brain regions compared to Allen’s CCFv3. For example, we found the granular layer of the olfactory bulb can be further divided into two layers. However, how this finer parcellation is related to the function of subregions remains to be determined.

With the 123 coronal brain sections from a single mouse brain, we have constructed the 3D transcriptome map of the whole mouse brain. This provided a unique opportunity for analyzing the gene module in a 3D manner. Although the mRNA distribution in the brain has been examined by ISH^2^, it is challenging to integrate all data as these data were collected in different animals. In contrast, we collected all data from a single mouse brain. We have taken thalamus as an example. Our analysis showed that few subregions of thalamus corresponded to one gene module, whereas most individual subregions in the thalamus gene corresponded to more than one gene module. This result suggests that transcriptomic characteristics only represent one aspect of the subregions. The parcellation of the subregion would need to combine different features. Limited by computation resources, we have not performed similar analysis on whole brain data. New efficient algorithms are required to process large amounts of dataset in the future.

### Limitation

We have characterized spatial distribution of all kinds of transcripts and various cell types in an unbiased manner. There are several limitations in our study. Firstly, due to the limitation in capture efficiency of gene transcripts, which has been a common issue for *in situ* based-spatial transcriptomic methods^4^, we could have missed some genes with low-expression level in our single cell spatial transcriptomic analysis. In fact, we identified 668 genes for one cell on average, which was not high enough for precise cell annotation. Therefore, we adopted Spatial-ID, an integrating transfer learning and spatial embedding strategy^27^, to perform cell type annotation for spatial transcriptomic datasets based on our snRNA-seq data with precise cell type annotation. Secondly, the number of brain sections and sampling time points for the developing mouse brain is limited, restricting the resolution and depth of our analysis. Thirdly, the coronal brain sections were collected at 100 µm-intervals for stereo-seq, which could have resulted in the missing of small nuclei and discontinuity of brain region reconstruction. Therefore, the sampling of the brain with smaller spatial intervals, or even profiling all sections across the whole brain, would be needed for a more precise 3D whole brain single cell transcriptome atlas in the future.

In summary, our study has constructed a 3D whole mouse brain single cell spatial transcriptome atlas, and mapped both the gene and cell cluster distribution across the mouse brain. We have identified 308 cell clusters with snRNA-seq and mapped their spatial distribution. We have discovered new region-specific genes including lncRNAs. We also showed that brain areas could be precisely defined based on the gene modules. Moreover, we found numerous TF regulons with spatial and temporal specificity. Thus, our construction of a whole mouse brain atlas with single cell resolution provides the basis for further exploring the function and development of the mouse brain.

## Supporting information

Table S1. Coronal sections information

Table S2. Cell clusters annotation

Table S3. Distribution of cell clusters ratio composition across brain areas

Table S4. Classification of regional-specific genes in brainstem

Table S5. Representative genes in spatial modules

Table S6. Spatiotemporal profile of gene expression in the developing mouse brains

Table S7. The representative lncRNA in different brain regions and spatial modules

Table S8. Gene modules enriched in brain regions during the developing mouse brains

Table S9. The correlation of divergence and convergence lncRNA-mRNA gene pairs across all spatial transcriptome samples

Table S10. Abbreviations of mouse brain areas

Movie S1. 3D demonstration of selected cell clusters

Movie S2. 3D demonstration of brainstem-specific neurons

Movie S3. 3D demonstration of brainstem-specific neuropeptides

Movie S4. 3D demonstration of region-specific lncRNAs

## ACKNOWLEDGEMENTS

This work was supported by the National Science and Technology Innovation 2030 Major Program (2021ZD0204400), Shanghai Municipal Science and Technology Major Project (Grant No.2018SHZDZX05), the National Natural Science Foundation of China (No. 3221003), the Scientific Instrument Developing Project of CAS (No. YJKYYQ20190052), National Key R&D Program of China (No. 2022YEF0203200, No.2022YFA1603604), Guangdong Provincial Key Laboratory of Genome Read and Write (2017B030301011) and New Cornerstone Science Foundation through the XPLORER PRIZE.

## AUTHOR CONTRIBUTIONS

H.Wang, J.Deng, Y.Zhong, J.Lin, Y.Chen, M.Xu, B.Ren, M.Cheng, Q.Yu, X.Song, Y.Lu, N.Yuan, S.Sun, Y.An, W.Ding, X.Sun, S.Zhang, Y.Dou, Y.Zhao, J.Xu, S.Wang, W.Ting, Y.Liu, M.Chen, C.Xie, B.Bo, X.Zhang, Z.Chen, J.Fang, S.Li, Y.Jiang, X.Tan, G.Zuo, H.Li, Y.Liu, J.Liu, M.Hao, J.Wang, J.Liu, H.Zhang, Y.Sheng, S.Yu, X.Jiang, G.Wang, C.Wang, X.Liu, X.Xu, H.Cao, H.Zheng, Y.Chen, H.Lu, Z.Yu, J.Zhang, B.Wang, Z.Wang, Q.Xie, S.Pan, C.Liu and C.Xu performed Stereo-seq and snRNA-seq experiments. L.Han, Z.Liu, Z.Jing, Y.Liu, H.Chang, J.Lei, Y.Peng, K.Wang, Y.Xu, W.Liu, Z.Wu, Q.Li, X.Shi, M.Zheng, H.Pan, R.Zhang, J.Wu, Y.Tang, Y.Wei, L.Han, Q.Zhu, H.Chen, D.Wang, Y.Bai, Y.Liang, M.Li, Y.Xie, Q.Tao, Y.Liu, H.Wen, Y.Yan, M.Han, X.Liao, H.Liu, N.Feng, K.Ma and T.Han analyzed the data. L.Cui, Y.Li, S.Liu, S.Liao, A.Chen, Q.Wu, J.Wang, Z.Liu, J.Mulder and H.Yang provided important advice for the project and comments on the manuscript. X.Wang, C.Li, J.Yao, X.Xu, L.Liu, Z.Shen, W.Wei and Y.Sun designed and supervised the study.

## RESOURCE AVAILABILITY

The processed data ready for exploration could be accessed via https://doi.org/10.12412/BSDC.1699433096.20001.

## Methods

### Tissue collection for Stereo-seq

Male mice (C57BL/6J) were used for the experiments. Left hemispheres of male mice (P1, P14, 11 wks) were collected. Animals were deeply anesthetized with 3% isoflurane and quickly perfused with 4°C artificial cerebrospinal fluid (ACSF, containing (in mM) NaCl 126, KCl 2.5, NaH_2_PO_4_ 1.25, MgCl_2_ 2, CaCl_2_ 2, NaHCO_3_ 26, and glucose 10, 300-305 mOsm) bubbled (with a mixture of 95% O_2_ and 5% CO_2_). After the dissection of the brain, the whole left hemisphere was obtained using the mouse brain sections (RWD, #68708). To prevent the formation of ice crystals during the tissue freezing, ACSF on the surface of brain blocks were dried with sterile gauze and brain blocks were transferred into OCT (4583#, Sakura) for 3 times to adequately displace the remaining ACSF. Subsequently, brain blocks were transferred to a self-made metal mold filled with OCT and quickly frozen with dry ice. After the freezing, brain blocks were stored in −80°C before the slicing. To minimize the RNA degradation, all solutions we used were prepared with diethyl pyrocarbonate (DEPC) (B501005-0005, Sangon Biotech) treated sterilized water (DEPC-H_2_O), and all instruments were washed with DEPC-H_2_O and RNase Zap (AM9780, Invitrogen). And time spent for the whole tissue collection process was constrained within 10 min.

### Tissue cryosection, section flattening and RNA quality control

Cryosection was performed to obtain one 10-µm section for Stereo-seq at 100-µm interval in coronal coordinate. The sections were carefully placed on pre-cooled Stereo-seq chips (−20°C) with a finger gently pressed under the chip which helps to reduce bubbles and wrinkles. RNA quality was examined, and only samples with RNA integrity number (RIN) value greater than or equal to 9 were used for analysis.

### Stereo-seq experiment procedure

The tissue section on the Stereo-seq chip (1 cm x 1 cm) was then baking at 37°C for 3 min and subsequently fixed in methanol (Sigma, 34860, precooled for 20 min at −20°C; 1 ml methanol was added in multi-orifice for each section) and incubated at −20°C for 30 min. Methanol was then dried out in a hood. Tissue section on the chip was then stained with ssDNA reagent (Invitrogen, Q10212) for 5 min and subsequently washed with 0.1x SSC buffer (Ambion, AM9770; containing 0.05 U/µl RNase inhibitor). Section images were captured using Zeiss Axio Scan Z1 microscope (at EGFP and DAPI wavelength, 10-ms exposure). Tissue sections were then permeated by incubating in 0.1% pepsin (Sigma, P7000, pepsin was prewarmed at 37°C for 3 min) at 37°C for 12 min (6 min for olfactory bulb) in 0.01 M HCl buffer (pH = 2) and then washed with 0.1x SSC buffer (containing 0.05 U/µl RNase inhibitor) to remove pepsin. In this step, RNAs were released from the permeated tissue and captured by the Stereo-seq chip. RNAs were then reverse transcribed for 1 hour at 42°C. After reverse transcription, tissue sections were washed with 0.1x SSC buffer and digested with tissue removal buffer (10 mM Tris-HCl, 25 mM EDTA, 100 mM NaCl, 0.5% SDS) at 37°C for 30 min, and then the chips were washed twice with 0.1x SSC buffer. The cDNA-containing chips were then subjected to Exonuclease I (NEB, M0293L) treatment for 3 hours at 55°C. The released cDNAs were collected and the chips were washed once with NF-water. The cDNAs were purified using 0.8x VAHTSTM DNA Clean Beads and then amplified with Hot Start DNA Polymerase (QIAGEN). The PCR reaction protocol was: first incubation at 95°C for 5 min, 15 cycles at 98°C for 20 s, 58°C for 20 s, 72°C for 3 min and a final incubation at 7°C for 5 min. The PCR products were then purified using 0.6x VAHTSTM DNA Clean Beads and were quantified by Qubit dsDNA HS assay kit (Invitrogen, Q32854).

### Stereo-Seq Library Construction and Sequencing

The resulting cDNA products were quantified using the Qubit™ dsDNA Assay Kit (Thermo, Q32854) after purification using VAHTS DNA Clean Beads (Vazyme, N411-03, 0.6×). 20 ng of cDNA products were fragmented using in-house Tn5 transposase at 55°C for 10 min, after which the reaction was stopped by the addition of 0.02% SDS. The fragmentation products were amplified with KAPA HiFi Hotstart Ready Mix, 0.3 µM Stereo-seq-Library-F primer and 0.3 µM Stereo-seq-Library-R. The reaction then ran: 1 cycle at 95°C for 5 min, 13 cycles at 98°C for 20 s, 58°C for 20 s, and 72°C for 30 s, and 1 cycle at 72°C for 5 min. The PCR products were then purified using VAHTS DNA Clean Beads (×0.6 and ×0.15). Finally, the library was sequenced on MGI DNBSEQ-Tx sequencer with sequencing length of 35 bp for read 1 and 100 bp for read 2.

### Processing of stereo-seq raw data

The stereo-seq libraries were sequenced by MGI DNBSEQ-Tx sequencer, and the fastq files generated were then processed with the SAW pipeline (https://github.com/BGIResearch/SAW). For read 1, coordination identity (CID) sequences were mapped to the designed coordinates of the *in situ* captured chip, allowing 1 base mismatch. Unique molecular identifiers (UMI) having either N bases or more than 2 bases with quality score less than 10 were filtered out. The associated CIDs and UMIs extracted from read 1 were appended to the read header of relative read 2. Then STAR^87^ was used to align retained read 2 to reference genome (mm10), and mapped reads with MAPQ >10 were kept. UMIs with the same CID and the same gene locus were collapsed, where one mismatch was allowed for sequencing and PCR errors. Finally, this information was used to generate an expression profile matrix containing coordinates.

### Cell segmentation

We harmonize the nucleic acid staining image and CID-containing expression matrix from the same section to perform spatial cell segmentation. Specifically, we utilized StereoCell (https://github.com/BGIResearch/StereoCell)^88^ to perform a three-step cell segmentation procedure. Firstly, staining images were registered with their corresponding expression matrices. Secondly, cell morphological segmentation was done based on nuclei-stained images. We employed median filtering to smooth the noise that may present in input staining images. Then, a modified U-Net based deep learning model^89^ was used to identify cell morphology from the image. Cells were filtered by their area, while shape and boundary was revised using opening operation (erode and dilate). Thirdly, after the basic nuclei mask was identified, a molecule labeling step is employed to label spatially detected UMIs to cells. The UMI that are contained within each nucleus are assigned directly to each cell. Then, to retrieve the UMIs in cytoplasm, a Gaussian mixture model was used to estimate the probability of each remaining UMI belonging to a given cell based on the initial nuclei segmentation, and label the UMIs with high confidence to the corresponding cells. Finally, we aggregated the UMIs that belong to the same cell, for each gene, and generated a cell-gene matrix for downstream analysis.

### Identification of anatomic regions

To identify anatomic regions, the expression profile matrix of both the coronal and sagittal mouse brain was divided into non-overlapping bins covering an area of 100 ×100 DNB. The UMIs were aggregated per gene within each bin. The resulting bins were further processed by Seurat (Hao et al., 2021). To be specific, we first performed quality control on the Bin100 spots of each brain section, filtering out bins with gene counts less than 200 and mitochondrial proportions higher than 5%. Then, we used the SCTransform function in the Seurat package to normalize the expression matrix of each Seurat object for each brain section separately. For the continuous sections of the whole mouse brain, we grouped the brain section data based on the interval from the appearance to the disappearance of major brain regions in the brain section (e.g. the 123 brain sections of mouse 1 were divided into 12 groups). We then merged the Seurat objects of brain sections in each group separately. We then used the RunPCA function to perform dimensionality reduction on the merged Seurat objects and performed unsupervised clustering on the normalized data using the FindNeighbors and FindClusters functions (parameters: reduction = 0.8, dims = 1:30). The clustering was performed using the Louvain clustering algorithm. Resulting clusters were annotated based on their transcriptomic marker genes identified by the FindAllMarker function of Seurat using default parameters. Anatomic annotation was further verified and compared with ABA (http://mouse.brain-map.org/).

### Image registration and brain region parcellation

The expression matrix of each section was used to generate total RNA images. The ssDNA and DAPI channel images scanned by ZEISS Z1 were spliced to extract the check lines, which were then registered with the corresponding check lines on the total RNA image so that the total RNA image, ssDNA image and DAPI channel image were aligned in the same coordinate system. Next, we identified the tilt angle and position corresponding to each section on the Allen 3D standard brain atlas. Marker points were then selected to perform cortical registration between the total RNA data and the standard brain atlas. Homemade CellPlot software was used to obtain parcellation of all cortical regions and subregions.

### 3D reconstruction of coronal mouse brain sections

2D sections registration: To register reconstructed brain sections into Allen Mouse Brain Common Coordinate Framework (CCFv3). We acquired the transformation matrix for each brain section based on 2D sections and CCFv3 reference and then applied this matrix to those reconstructed cells and genes in this brain. First, we use 3D section software to register images by pairs of manually selected fiducial points so that the outlines are as aligned as possible. In this process, we will get the transformation matrix for each brain section based on 2D sections. Next, the resultant transformation matrix was applied to the reconstructed cells and genes in the brain.

### Single-nucleus brain areas sampling

Mouse was anesthetic with the mixture of Loratadine Tablets (50 mg/kg) and Xylazine Hydrochloride (0.1 mg/kg), and perfused with ice-cold artificial cerebral spinal fluid (ACSF) containing (in mM) NaCl 126, KCl 2.5, NaH_2_PO_4_ 1.25, MgCl_2_ 2, CaCl_2_ 2, NaHCO_3_ 26, and glucose 10 (300-305 mOsm). The mouse brain was quickly dissected and transferred to the dish (diameter: 6 cm) that contained bubbled ice-cold ACSF. Then, the brain was divided into two parts along the coronal axis, the dividing site dependent on the tissue that we need to correct. After that, the part that we needed was attached to the platform of the cutting system (Leica VT1200S) and the platform was attached to the cutting chamber which was filled with bubbled ice-cold ACSF. Coronal slices (500 µm) were prepared at the speed of 0.2 mm/s with blade vibration amplitude of 0.8 mm. Immediately after the section, brain slices were transferred to the dissection chamber (containing bubbled ice-cold ACSF). Then, target brain areas were carefully dissected according to the allen-atlas and collected to responding tubes that stored on the dry-ice. After the collection of all brain areas of the mouse, tubes were put into the liquid nitrogen for 3 min and then transferred into the dry-ice. The duration should be less than 10 min from the time point of perfusion to that of tubes into liquid nitrogen. After the daily collection of all tissues, the boxes that contain dry-ice and tubes were stored in the −80 refrigerator before being sent out for sequencing.

### Single-nucleus suspension preparation

Similar to previously described^90^, frozen brain tissues were rapidly transferred to a 1-ml Dounce homogenizer (TIANDZ) and immediately homogenized in a 1 ml homogenization buffer. After filtered with a cell strainer into a 1.5-ml tube (Eppendorf), solution in the tube was subsequently centrifuged at 500 g for 5 min at 4°C to pellet the nuclei. Then, the pelleted nuclei were diluted in a resuspension buffer at 1000 nuclei/µl for library preparation.

### Single-nucleus library preparation and sequencing

Single-nucleus RNA sequencing libraries were prepared using DNBelab C Series High-throughput Single-Cell RNA Library Preparation Kit (MGI, #940-000047-00). In brief, single-nucleus suspensions were processed as follows: droplet generation, emulsion breakage, beads collection, reverse transcription and cDNA amplification to generate barcode libraries. Indexed libraries were prepared according to the manufacturer’s protocol (MGI, 1000021082). Qubit ssDNA Assay Kit (Thermo Fisher Scientific, Q10212) was used to quantify the constructed libraries. The library was sequenced on DNBSEQ-T1 or DNBSEQ-T7 sequencer with following read length: 41-bp length for read 1, 100-bp read length for read 2 and 10-bp read length for sample index.

### Single-nucleus data generation

Raw data processing. Raw sequencing reads were processed with filtering, demultiplexing barcode processing and 3’ UMI counting using the DNBelabC Series HT scRNA analysis Software Suite (v1.0.0) set with default parameters (https://github.com/MGI-tech-bioinformatics/DNBelab_C_Series_HT_scRNA-analysis-software/tree/version1.0). PISA software (https://github.com/shiquan/PISA) was applied to parse raw reads into FASTQ format according to library structure and cell barcode information. Processed reads were then aligned to GRCm38.p3 (mm10) mouse genome with STAR (v2.7.4a) and sorted by sambamba (v0.7.0). Due to the large amount of unspliced pre-mRNA in mouse brain cell nucleus, a custom ‘pre-mRNA’ reference was created for alignment of count reads to exons as well as to introns. Accordingly, each gene’s transcript in snRNA-seq was counted by exon and intron reads together. Finally, a nucleus-gene metric was generated, which was then adjusted by SoupX (v1.5.21) to reduce ambient RNA noise.

### Single-nucleus data filtering

In all single-nucleus sequencing libraries, we removed libraries with median gene count ≤ 800 and selected the remaining 136 libraries as high-quality libraries. Nuclei were further filtered based on criteria including median gene count > 800, mitochondrial gene content < 5%, and a ratio of UMI counts to gene counts > 1.2. The expression data were then subjected to SCTransform normalization in the Seurat package (v4.3.0). We performed unsupervised clustering on the normalized data using the FindNeighbors and FindClusters functions (parameters: dims = 1:50). Based on the marker genes of each group, the clustering results were divided into six major cell classes: *Neurons*, *Astrocytes*, *Oligodendrocytes*, *Microglia*, *Ventricular cells*, and *Vascular cells*. Nuclei that could not be clearly classified into these six major types were classified as *Uncertain*.

When dividing nuclei into major classes, it was found that some nuclei existed at the boundaries between classes and were classified as the uncertain class without clear marker genes. Therefore, we trained a random forest model to predict the major nuclei classes in order to remove the uncertain nuclei. First, we downsampled each nuclei class and extracted 5,000 nuclei (up to 80% of the type) as the training set. The FindAllMarkers function in the Seurat package was used to identify marker genes for each class in the training set, and further filtered genes with corrected *p* values (p_val_adj) < 0.01 and average log_2_ fold change (avg_log_2_FC) > 1 as feature genes for the random forest model. Using the SCTransform normalized values as gene-expression data, the randomForest function in the randomForest package (v4.7.1.1) was used to build the model based on the training set (parameter: ntree = 500). Then, the predict function was used to apply the trained model to all single-nucleus data. The model evaluated the probability of the major classes for each nucleus, and we assigned each nucleus to the class with the highest probability. Nuclei with highest probability < 0.9 were discarded. After the iterative clustering and annotation that introduced below, clusters with unclear marker expression were removed. 378,287 nuclei were kept after all filtering processes and used in further analysis.

### Single-nucleus iterative clustering and classification

The single nucleus data was first normalized using SCTransform and reduced the dimension using RunPCA. We used the modified RunHarmony function in the harmony package (v0.1.0) to integrate data from different batches. We modified the kmeans clustering algorithm used in the package by adding a new parameter kmeans.algorithm, which replaces the default Hartigan-Wong algorithm with the Lloyd algorithm. We also added two parameters, kmeans.iter.max and kmeans.nstart, to make it applicable to larger datasets (parameters: assay.use =“SCT”, reduction = “pca”, dims.use = 1:50, theta = 2, max.iter.harmony = 10, max.iter.cluster = 20, kmeans.algorithm = “Lloyd”, kmeans.iter.max = 1000, kmeans.nstart = 20, epsilon.cluster = −Inf, epsilon.harmony = −Inf).

To further subdivide cell types, we followed the iterative clustering method for single-cell sequencing data in the scrattch.hicat package (https://github.com/AllenInstitute/scrattch.hicat), while using the batch correction method in harmony to correct the batch effects in different batches during each iteration. This allowed us to reduce batch effects while clustering cell clusters in as much detail as possible. The specific steps are shown as follows:

(1) First, we defined a clustering function, OnestepCluster, which takes the single cell data in Seurat object format and performs SCTransform normalization, RunPCA dimension reduction, and RunHarmony batch correction. The clustering results are checked using the MergeCluster function in step (2) to merge the clusters without distinguishable gene expression differences.

(2) The MergeCluster function calculates the Pearson correlation coefficient between any two clusters. Clusters with fewer than 20 cells will be merged directly with the other cluster with the highest Pearson correlation coefficient, and then the Pearson correlation coefficient is recalculated for all clusters. Next, we calculated the gene expression differences between the cluster and the other top 3 clusters with the highest Pearson correlation coefficient. The FindMarkers function is used to calculate the differentially expressed genes G between any two clusters. To shorten the computation time, each cluster is downsampled to 500 cells for calculation. The differential gene score S is calculated based on the following formula:

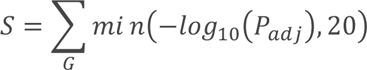

Among them, differentially expressed gene G requires corrected *p* values < 0.01 and average fold change > 1, with at least 50% of cells expressing the gene in the highly expressed cluster, and the expression proportion of the gene in the lowly expressed cluster is more than 50% lower than the highly expressed clusters. If the number of differentially expressed genes G is less than 5 or the differential gene score S is less than 200, the two clusters will be merged. All calculations will be repeated until the differential gene score S between any two clusters larger than 200.

(3) We defined the clustering recursive function IterCluster, which combines the OnestepCluster and MergeCluster in step (1-2). If all clusters are merged in the result, the recursion will be terminated because it cannot be further split. If there are more than one clusters in the result, the clusters with more than 50 cells are selected and the IterCluster function is performed on this cluster again. The process continues until all clusters cannot be further splitted, and all clustering results are summarized and returned.

At the beginning, we used the results of the six major nuclei classes as the start point. For each major class, the IterCluster recursive function is executed, and eventually, all clusters splitting results are returned. We annotated the clusters based on the marker genes, and removed some groups with unclear marker genes. In the end, we obtained 308 clusters covering 378,287 high quality nuclei.

### Single-nucleus cluster dendrogram construction

Dendrogram of 308 nuclei clusters was constructed based on the marker gene expression. First, the FindAllMarkers function in Seurat software package was used to identify marker genes for all cell types. Only genes with adjusted *p* values (p_val_adj) < 0.05 and avg_log_2_FC > 0 were kept. For each nuclei cluster, the top 20 genes with the largest avg_log_2_FC were selected. Marker genes with expression in less than 20% of nuclei in the corresponding cluster were filtered out. Next, the mouse whole-brain snRNA-seq data were divided into two groups: neuronal and non-neuronal nuclei. For each group, the AverageExpression function was used to calculate the average expression values of marker genes for each nuclei cluster. The pvclust function in the pvclust package (v2.2.0) was used to construct a classification dendrogram, based on the Spearman correlation distances between clusters. Finally, the merge function was used to combine two dendrograms together.

### Independence test of nuclei clusters using random forest model

We tested the independence of each cluster in the iterative clustering results to ensure there are sufficient differences between nuclei clusters. We used the randomForest function in the randomForest package to build a random forest model for all 308 clusters. We first used the FindAllMarkers function in the Seurat package to find the marker genes for each cluster, Only the genes with an adjusted *p* value (p_val_adj) < 0.05 and an avg_log_2_FC > 0. For each nuclei cluster, we selected the top 50 genes with the highest avg_log_2_FC and used them as the feature genes for the random forest model. Then, the single-nucleus data was downsampled, and 200 nuclei were selected for each nuclei cluster. Using the SCTransform-normalized values as expression data, we built the random forest model using the randomForest function (parameters: ntree = 1000). The confusion matrix was calculated using the out-of-bag (OOB) data of the model, Prediction accuracy of the confusion matrix data was used as the standard for evaluating independence clusters.

### Single-nucleus data co-clustering with published dataset

We used two public scRNA-seq datasets as references. One of which is the mouse whole nervous system scRNA-seq dataset^3^, we removed cell types from the spinal cord and peripheral nervous system since these cells were not included in our snRNA-seq dataset. For the other public dataset^26^, which only contains single-cell data from the cortex and hippocampus structures, we extracted cell types belonging to these two regions from our dataset and discarded clusters with cell counts < 20 for comparison purposes.

We first downsampled 200 cells for each cell type in each dataset separately. Using the SCTransform function in the Seurat package (v4.3.0), we normalized each dataset (parameters: variable.features.n = 5000, method = “glmGamPoi”) and reduced the data using the RunPCA and RunUMAP functions (parameters: dims = 1:50). We used the FindAllMarkers function to find marker genes for each cell type and only kept genes with a corrected *p* values (p_val_adj) < 0.05 and an avg_log_2_FC > 0. We then summarized the top 50 genes with the highest avg_log_2_FC for each cell type as the marker gene set for that dataset. Using the FindIntegrationAnchors and IntegrateData functions, we integrated our dataset with any one of the publicly available datasets, using the intersection of marker gene sets as integration anchors. We then reduced the integrated data again using RunPCA and RunUMAP (parameters: dims = 1:100), and performed clustering using FindNeighbors (parameters: reduction = “pca”, dims = 1:100) and FindClusters (parameters: resolution = 3, n.start = 10, algorithm = 1).

For the integrated clustering results, we calculated the proportion R of a cell type x in our dataset that clusters with a cell type y in other public dataset using the following formula:

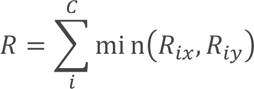

where C represents all clusters generated after re-clustering the integrated dataset, i represents any cluster in C, Rix represents the proportion of cell type x in cluster i among all cell types of x, and Riy represents the proportion of cell type y in cluster i among all cell types of y.

### Stereo-seq cell type transfer

Due to the intricate diversity of neuron types throughout the entire brain and the wide-ranging spatial transcriptomic regions, we partitioned both the snRNA-seq data and Stereo-seq data into six distinct, mutually independent regions (cerebral cortex, cerebral nuclei, interbrain, midbrain, hindbrain, cerebellum). When transferring cell types to a specific region within the Stereo-seq data, we selected the snRNA-seq neuron clusters from the corresponding region as the reference set. Clusters comprising less than 20 nuclei were excluded. Considering that non-neuronal cells exhibit relatively lower regional specificity in their distribution across the entire brain, we utilized all non-neuronal cells from the complete mouse brain snRNA-seq dataset as the reference set. Each region in Stereo-seq data was transferred using the corresponding neuronal and all non-neuronal cell clusters. In addition, the Stereo-seq data also included fiber tracks and ventricular regions, which are not primary structural regions of the mouse brain and predominantly comprise non-neuronal cells. Consequently, for these regions, we exclusively employed non-neuronal cells as the reference set for cell type prediction. We defined the cells with more than 100 genes and the percentage of mitochondrial genes less than 15%. Prior to cell type transfer, each cell cluster in the snRNA-seq data was downsampled to 1000 nuclei. We computed the marker genes for each cluster, selecting the top 300 genes (sorted by fold changes, excluding genes associated with mitochondria or ribosomes), which were subsequently utilized in the subsequent training phase.

We used Spatial-ID, a novel cell type mapping algorithm^27^, to annotate single cells in Stereo-seq data with cell types classified by snRNA-seq analysis. Cells in Stereo-seq data with detected genes less than 100 and percentage of mitochondrial genes larger than 15% were removed in the transfer process. The annotation procedure mainly consisted of two stages. In the first stage, we trained a four-layer deep neural network (DNN) with the snRNA-seq data. The DNN model included two hidden layers of 2048 and 1024 nodes and was trained by a Cross Entropy loss with learning rate = 3×10^-4^, weight decay = 1×10^-6^. The trained DNN predicted the initial probabilities of defined cell types for single cells on the Stereo-seq data. In the second stage, we applied a graph convolution network (GCN) including two autoencoders and a classifier to integrate gene expression, spatial neighborhood information and cell type instruction generated in the previous stage to refine the probabilities. Spatial neighborhood information of the Stereo-seq data was encapsulated into an adjacency matrix of cells with normalized Euclidean distances as values for non-zero elements. The GCN model took the gene expression matrix, the distance-weighted adjacency matrix and the initial probabilities as inputs and produced the final probabilities after 200 epochs of training with learning rate = 1×10^-2^, weight decay = 1×10^-4^. The two autoencoders of the GCN model were trained via a self-supervised learning strategy with reconstruction losses and the classifier was trained by a Cross Entropy loss. Spatial-ID used in this study was implemented with PyTorch (v1.13.1) and PyTorch Geometric (v2.1.0) packages in Python (v3.8.10).

### Analysis of cell type distribution density across different brain regions

We calculated the density distribution of cell types across 66 brain regions using coronal slices. Density is derived by dividing the count of specific cell types within each region by the corresponding region’s area. The cell type count is determined through the analysis of coronal slices, while the region’s area is computed by tracing lines using total RNA and registered ssDNA images of the coronal slice. Subsequently, utilizing summary statistics of density values from various slices within the same region, each data point signifies the mean density across all brain slices within that region.

### Analysis of cell type distribution density along anterior-posterior axis

In Figure 2B, the x-axis illustrates the distribution of distances along the anterior-posterior axis. The y-axis represents ten distinct cell types specific to regions, while the z-axis depicts the relative counts of cell types within the coronal section. Relative counts were determined by dividing the cell count within the coronal section by the total cell count in that section, and this data was obtained from the transfer results. Each data point signifies the relative count of a specific cell type within a coronal section at a distinct location along the anterior-posterior (AP) axis.

### Regulon analysis

The regulon activity analysis of transcript factors was performed using standard pySCENIC pipeline^91^. Transcriptomic profiles of previously divided bins (bin100) were used as input. The GENIE3 algorithm^92^ was used to reconstruct the co-expressed gene network for each transcript factor. This network was then analyzed and filtered according to cisTarget database (https://resources.aertslab.org/cistarget/), resulting in regulons consisting of each TF and its potential targets. The activity of a regulon within each bin was calculated by AUCell using default parameters and threshold, and then mapped to physical space for visualization. We aggregated potential target genes sharing the same TF across different developmental time points by using the *union* function. Aggregated regulons were subjected to AUCell again to re-calculate comparable activity scores across each developmental stage. The regulon activity of each anatomic region was defined as the mean activity of all bins belonging to the region.

### Identification of spatial gene modules

Spatially co-expressed genes and co-activated regulons are identified and grouped into modules using Hotspot^32^. Top 5000 highly variable genes or all identified regulons were used as input. Gene expression in each bin100 was scaled by size factor and log transformed. Then the spatial autocorrelation score was calculated using *compute_autocorrelations* function. Significantly auto-correlated genes or regulons with *p* value less than 0.05 were kept and further grouped into modules using the *create_modules* function with min_gene_threshold = 10 and fdr_threshold = 0.05 (for the regulon: min_gene_threshold = 5, C > 0.15 and fdr_threshold = 0.05, for genes in developmental sections: min_gene_threshold = 20 and fdr_threshold = 0.05). Identified gene or regulon clusters were annotated to related anatomic brain regions according to their spatial localization and contents of genes or TFs.

We identified gene modules with spatial specificity through gene module enrichment analysis of 123 brain sections from mouse #1. To identify gene modules with regional specificity among these modules, we calculated the enrichment scores of gene modules in bin100 spots from different brain regions of each brain section. For a given gene module in a particular brain section, we calculated the significance of the difference in gene module enrichment scores for bin100 spots across brain regions. If the enrichment score of a gene module in a specific brain region was significantly higher than that in other brain regions (p_val_adj < 0.01), we considered this gene module to be region-specific for that brain region. Due to the fact that computer-based methods cannot accurately annotate gene modules that cover multiple brain regions, we combined computational and manual methods to determine the brain region specificity of each gene module and annotate the gene modules.

### Gene ontology analysis and gene set enrichment scoring

Gene ontology (GO) analysis was performed on identified modules using clusterProfiler^93^. Gene members of each gene module were used as input. For TF regulon clusters, corresponding TFs and top 10 target genes of each regulon member were used. GO enrichment score and significance for biological process (BP), Molecular Function (MF) and Cellular Component (CC) were calculated with the compareCluster function using the *org.Mm.eg.db* database with default parameters. BP, MF, CC with Benjamini-Hochber-adjusted *p* values less than 0.05 were considered to be enriched in corresponding modules. For manually chosen developmental related process, we extracted the gene sets from Mouse Genome Informatics website (http://www.informatics.jax.org/vocab/gene_ontology), and calculated enrichment scores for these gene set within each bin100, using the AddModuleScore function in Seurat^94^ package. The results were then visualized in groups by brain sections, using the ggplot2 package.

### Spatial expression pattern analysis

To quantify and analyze the spatial distribution of transcriptomic features, cortex region of each developmental mouse brain section was digitized and divided into conformal layers or columns along the dorsal-ventral and rostral-caudal axis using the Spateo package digitization pipeline^41^. Concretely, for each brain section, cortex-corresponding clusters were subjected to the extract_cluster_contours function, and converted into a closed contour line indicating the boundary of cortex region. We arbitrarily divided and labeled the boundary of cortex into 4 segments, indicating the dorsal, ventral, rostral and caudal side of cortex, respectively, which were further used by digitize function to generate relative dorsal-ventral and rostral-caudal coordinates for each bin100 within cortex. We then performed differential analysis on gene expression or regulon activities distribution along the dorsal-ventral axis to identify laminar peak shifting in the developmental process as described in Figure 5F. Generalized linear models are applied on expression distribution of the rostral-caudal axis to detect significant gradient patterns, using the glm_degs function in Dynamo package^95^ with default parameters. Genes with Benjamini-Hochber-adjusted *p* values greater than 0.05 or expressed in less than 25 percent of bins in cortex were filtered out. Then, min-max normalized expressions of each significant gradient gene were further fitted using sklearn.linear_model. LinearRegression^96^ to get a normalized gradient coefficient and a coefficient of determination, where genes with coefficient of determination greater than 0.05 were shown in Figure 5G and Figure S6C.

### Identification of region-specific genes

For identification of region-specific genes, the gene density of each region was calculated as nCounts/region size, then the fold change was calculated as the highest region density/median density of all regions. Region-specific genes were defined as fold change > 1.5 and density > 25 per mm^2^. Finally, some genes were added or deleted based on manual screening by checking expression images of individual genes obtained by spatial transcriptomic experiment. Layer specific genes were calculated similarly except the cortex was manually divided into layer 1-6.

### Identification of convergent gene pair and divergent gene pair

We divided lncRNA-mRNA gene pairs into divergent and convergent groups based on the distance between the transcription start sites of lncRNA and mRNA on the genome. In a gene pair, lncRNA and mRNA are close or overlap and need to be on different DNA strands. When the transcription start site (TSS) distance between lncRNA and mRNA is greater than −1000 nt and less than 10000 nt, we define it as a divergent gene pair (TSS distance = TSS in plus strand – TSS in minus strand). When the TSS distance between lncRNA and mRNA is less than −1000 nt and lncRNA overlaps with mRNA, we define it as a convergent gene pair.

### Gene pair correlation in different brain regions

To calculate the correlation of lncRNA-mRNA expression in different brain regions of mice, we combined the gene expression levels in the same brain region, and then calculated the CPM value of each gene in each section. Then filter out low expression values (lncRNA + mRNA >1), and calculate the lncRNA-mRNA Pearson correlation coefficient.

### Calculation of gene pairs with differential expression within distinct cell types

To calculate the differential expression of lncRNA-mRNA in different cell types, we calculated the average expression of each gene in different cell types. Then calculate the fold change and *p* values (p_val_adj) of lncRNA and mRNA in the two cell types (FC = log_2_ (lncRNA/mRNA)). When the *p* values (p_val_adj) is less than 0.01, the absolute value of fold change is greater than 1, and the signs of FC in the two cell types are opposite, we consider that lncRNA and mRNA are antagonistic expressed in these two cell types.

## Supplementary figures

**Figure S1.**
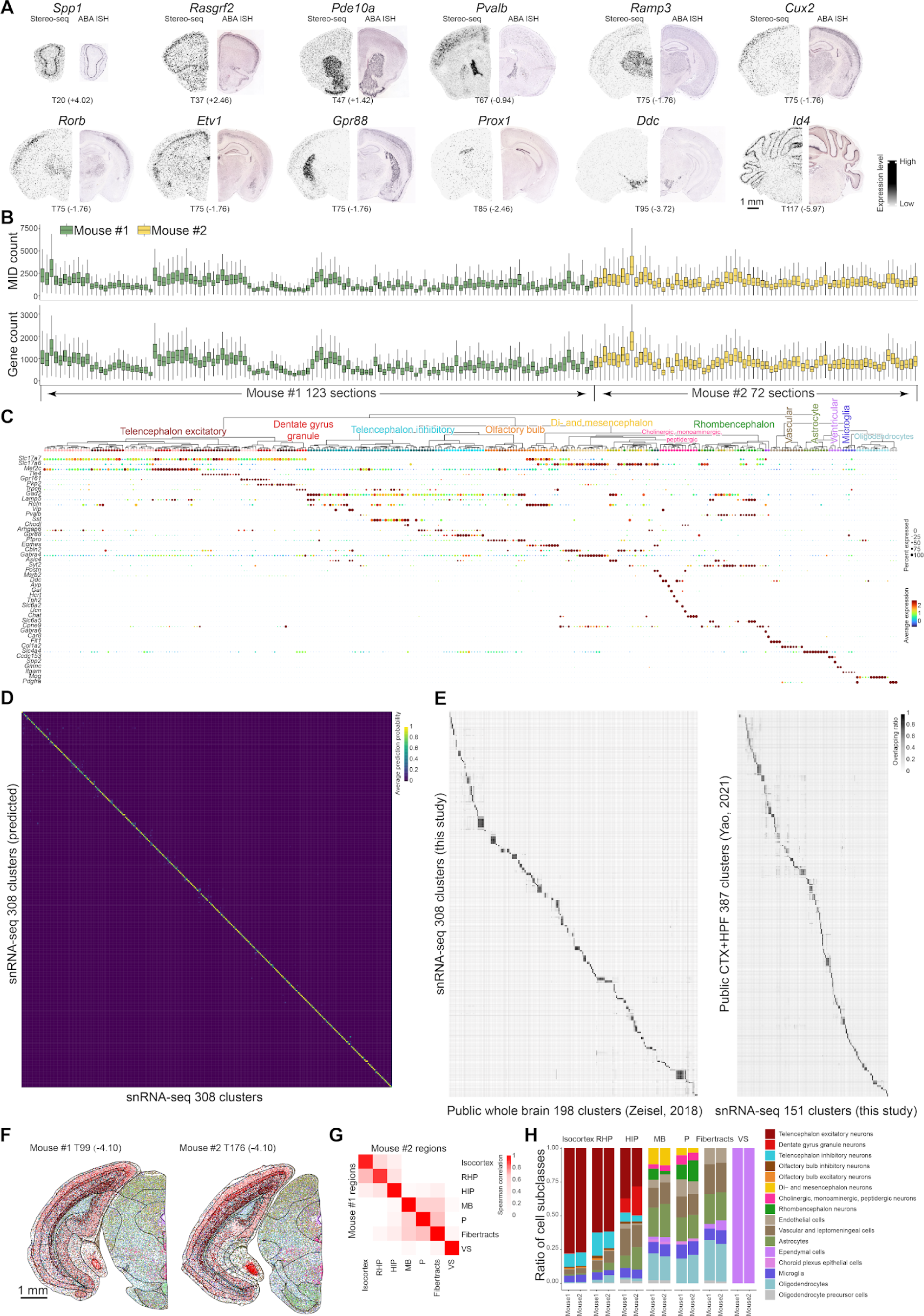
Benchmark of snRNA-seq and Stereo-seq data. **(A)**The distribution comparison of example genes between Stereo-seq spatial transcriptome (left side) and Allen Brain Atlas *in situ* hybridization (ABA ISH) images (right side) (section No. and bregma coordinates shown, unit mm). Scale bar, 1 mm. **(B)** The number of molecular identifiers (MIDs) and genes in cells from Stereo-seq data. Each box indicated one coronal section in two mouse brains Stereo-seq data (green for mouse1, yellow for mouse2). **(C)** The expression of marker genes of 308 cell clusters from snRNA-seq data. **(D)** Heatmap of the confusion matrix of random forest classifiers for 308 cell clusters from snRNA-seq data. **(E)** Heatmap of comparisons of snRNA-seq cell clusters with two published mouse brain single-cell datasets. **(F-H)** Replicates of two coronal sections from two mice. **(F)** Overall distribution of cell clusters in two sections. Scale bar, 1 mm. The cell clusters from the two mice are depicted using the same color scheme. **(G)** Heatmap of spearman correlation of the number of cells within cell clusters across different regions on the two sections. Cell clusters with fewer than 10 cells were excluded from the analysis. **(H)** The ratio comparisons of cell subclasses between two sections.

**Figure S2.**
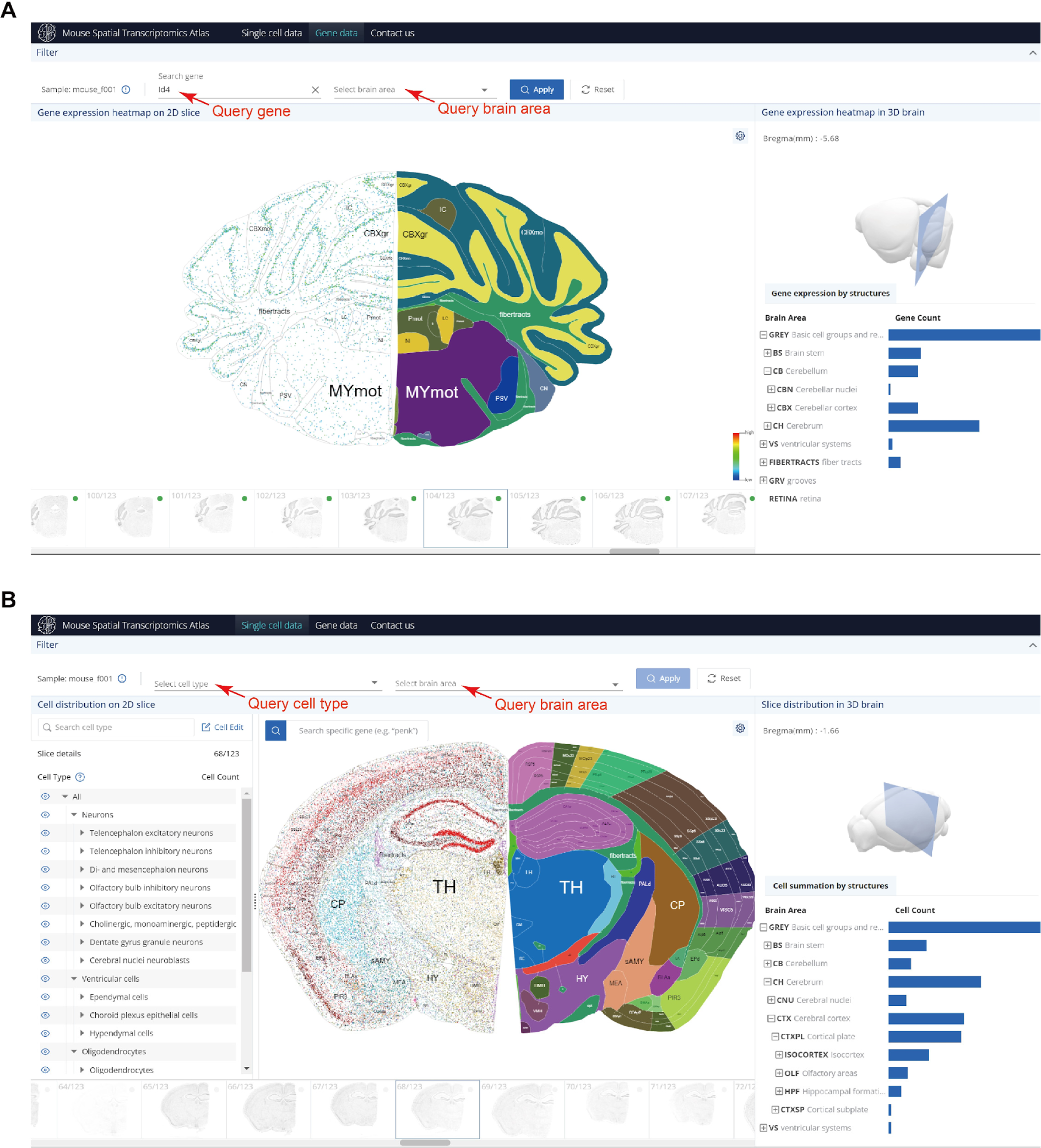
Interactive website depicting gene and cell distributions across the mouse brain. **(A)**Gene expression query page. Users can search for specific gene expressions in selected brain areas. **(B)** Cell type distribution query page. Users can explore specific cell type distributions in selected brain areas.

**Figure S3.**
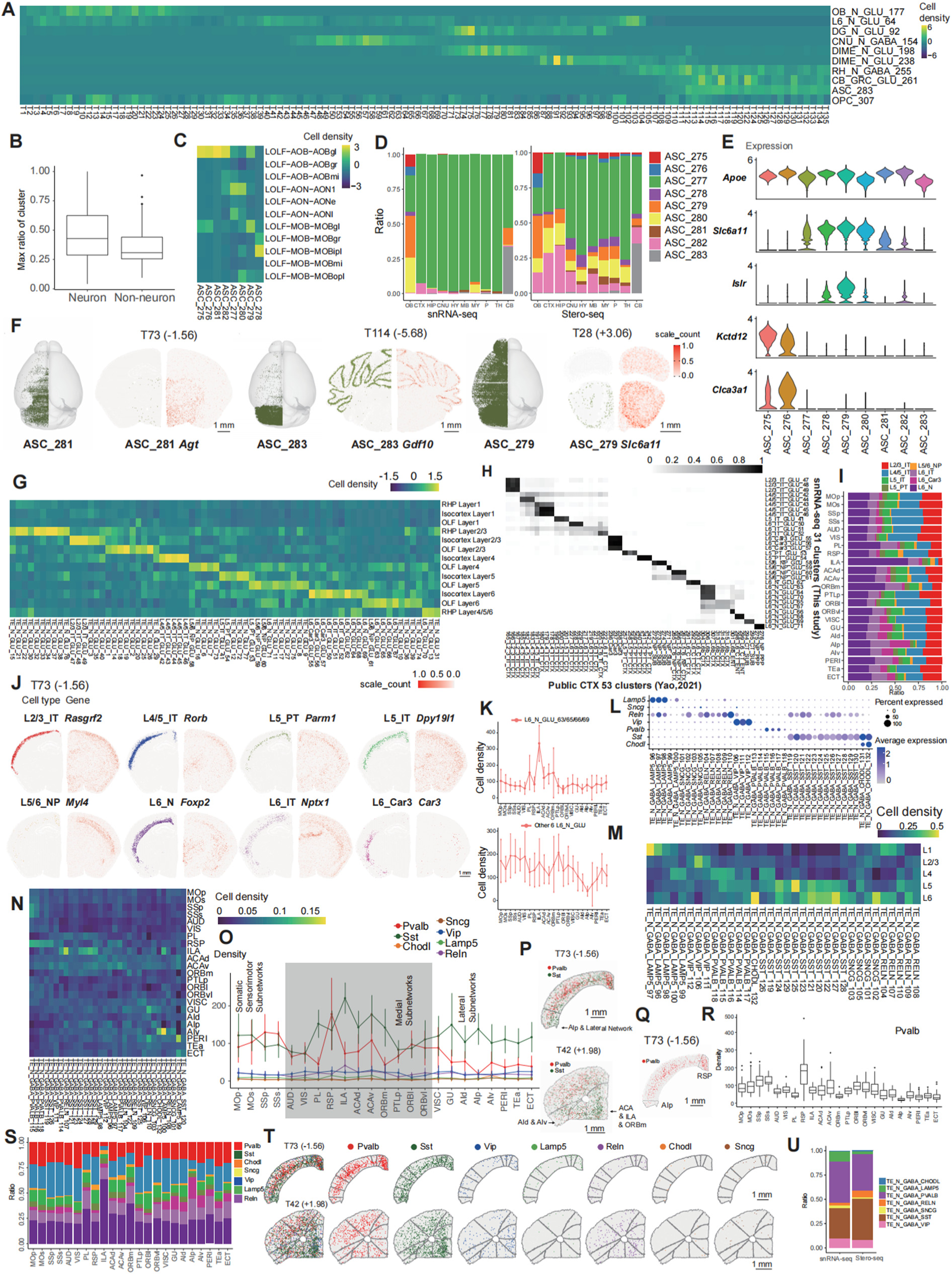
Distribution of excitatory neurons, inhibitory neurons, and non-neurons in the mouse brain. **(A)**A heatmap showing the distribution of ten representative cell clusters in coronal sections, calculated by relative numbers (number of representative cell clusters per section/ total number of cells per section). **(B)** Boxplot of the maximum of the cell ratio of neurons (262 clusters) and non-neurons (46 clusters) in 66 brain areas. The cell ratio of cluster x in area y is calculated by the cell number of cluster x in area y divided by total cell number of the cluster x. The maximum cell ratio of cluster x among all brain areas was used in the boxplot. **(C)** Heatmap of the distribution of 8 astrocyte cell clusters across different areas of the olfactory bulb. **(D)** Barplot of the proportion of 9 astrocyte cell clusters in different brain regions along anterior to posterior between snRNA-seq and Stereo-seq data. **(E)** Violin plot of the expression levels of *Apoe*, *Alc6a11*, *Islr*, *Kctd12*, *Clca3a1* in 9 astrocytes clusters in snRNA-seq data. **(F)** Spatial distribution of 3 representative astrocytes cell clusters (ASC_281, ASC_283, ASC_279). The demonstrations of their corresponding 3D spatial visualization, cell clusters distribution, marker gene expression on coronal sections are shown. Scale bar, 1 mm. **(G)** Heatmap of the distribution of all excitatory neurons clusters across layers of the OLF, isocortex and RHP. **(H)** Heatmap of the comparison of glutamatergic cell clusters between our data (31 cell clusters of glutamatergic excitatory neurons) and public data. **(I)** Barplot of the proportion of 8 excitatory cell groups in different areas of the isocortex. **(J)** Distribution of 8 excitatory cell groups (left) and expression of their marker genes (right). Scale bar, 1 mm. **(K)** Density distribution of ILA region-specific layer 6 neurons (L6_N_GLU_63, L6_N_GLU_65, L6_N_GLU_66, L6_N_GLU_69) and other layer 6 neurons in different areas of isocortex. Error bars were estimated across all cortical sections (mean ± SD). **(L)** Bubble plot of the expression levels of marker genes in 34 cell clusters of inhibitory neurons. **(M)** Heatmap of the distribution of 34 inhibitory neuron cell clusters across layers of isocortex. **(N)** Heatmap of the scaled density distribution of cortical inhibitory neuron cell clusters across cortical areas. **(O)** Cell density of the 8 inhibitory cell groups across anatomical regions of the isocortex arranged in 3 subnetworks based on their anatomical connectivity (Zingg et al., 2014), error bars were estimated across all cortical sections (mean ± SD). **(P)** The distribution of inhibitory neurons in coronal sections. Pvalb neurons are in red, Sst neurons are in green, and other cells are in gray. Scale bar, 1 mm. **(Q)** Pvalb neurons distribution in cortical regions of section T73. Scale bar, 1 mm. **(R)** Boxplot of density distribution of Pvalb neurons in different areas of the isocortex. **(S)** Barplot of the proportion of 7 inhibitory cell groups in different areas of the isocortex. **(T)** The distribution of inhibitory neurons in cortical areas (section T42 and T73). Scale bar, 1 mm. **(U)** Barplot of the cell ratio of inhibitory neurons in snRNA-seq and Stereo-seq data.

**Figure S4.**
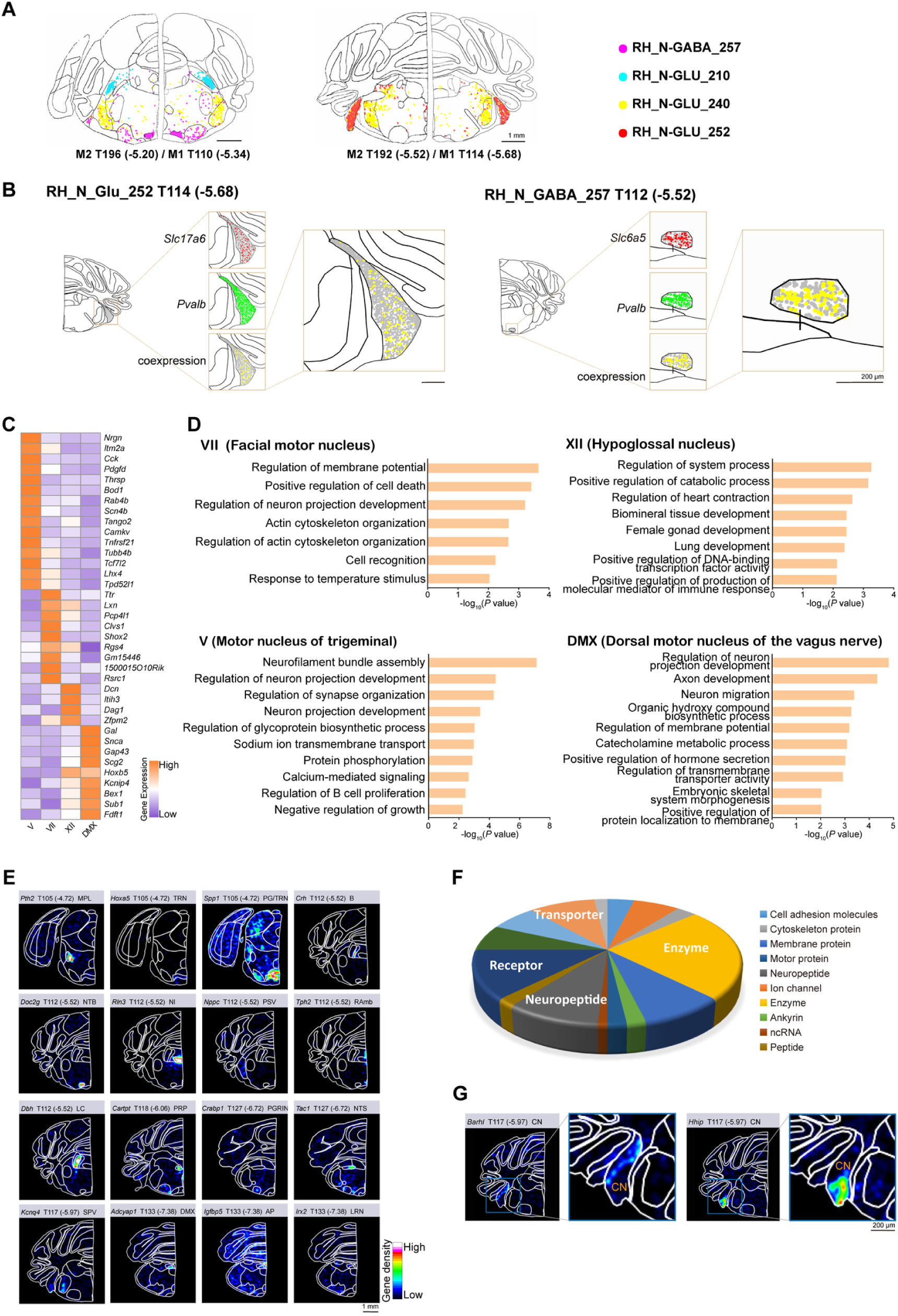
Region-specific cell clusters and genes in the brainstem. **(A)**Spatial visualization of 4 nuclei-specific cell clusters in representative brain sections of mouse #1 and mouse #2. The number in the parentheses on the bottom of each brain section represents the distance from bregma point. Scale bar, 1 mm. **(B)** Spatial visualization of the expression of *Slc17a6* and *Pvalb* in RH_N_GLU_252 in the cochlear nuclei (CN), and the expression of *Slc6a5* and *Pvalb* in RH_N_GABA_257 in the nucleus of the trapezoid body (NTB). The number in the parentheses on the top of the brain section represents the distance from the bregma point. Scale bar, 200 µm. **(C)** Heatmap showing the expression of 4 motor nucleus-specific genes in mouse #2. **(D)** Gene ontology enrichment analysis of the differentially expressed genes of the 4 motor nuclei. **(E)** Spatial visualization of representative region-specific genes in the brainstem. The number in the parentheses on the bottom of each brain section represents the distance from bregma point. Scale bar, 1 mm. **(F)** Pie chart shows the category of region-specific genes in the brainstem. **(G)** Representative subregion-specific genes in the CN. The number in the parentheses on the top of each brain section represents the distance from bregma point. Scale bar, 200 µm.

**Figure S5.**
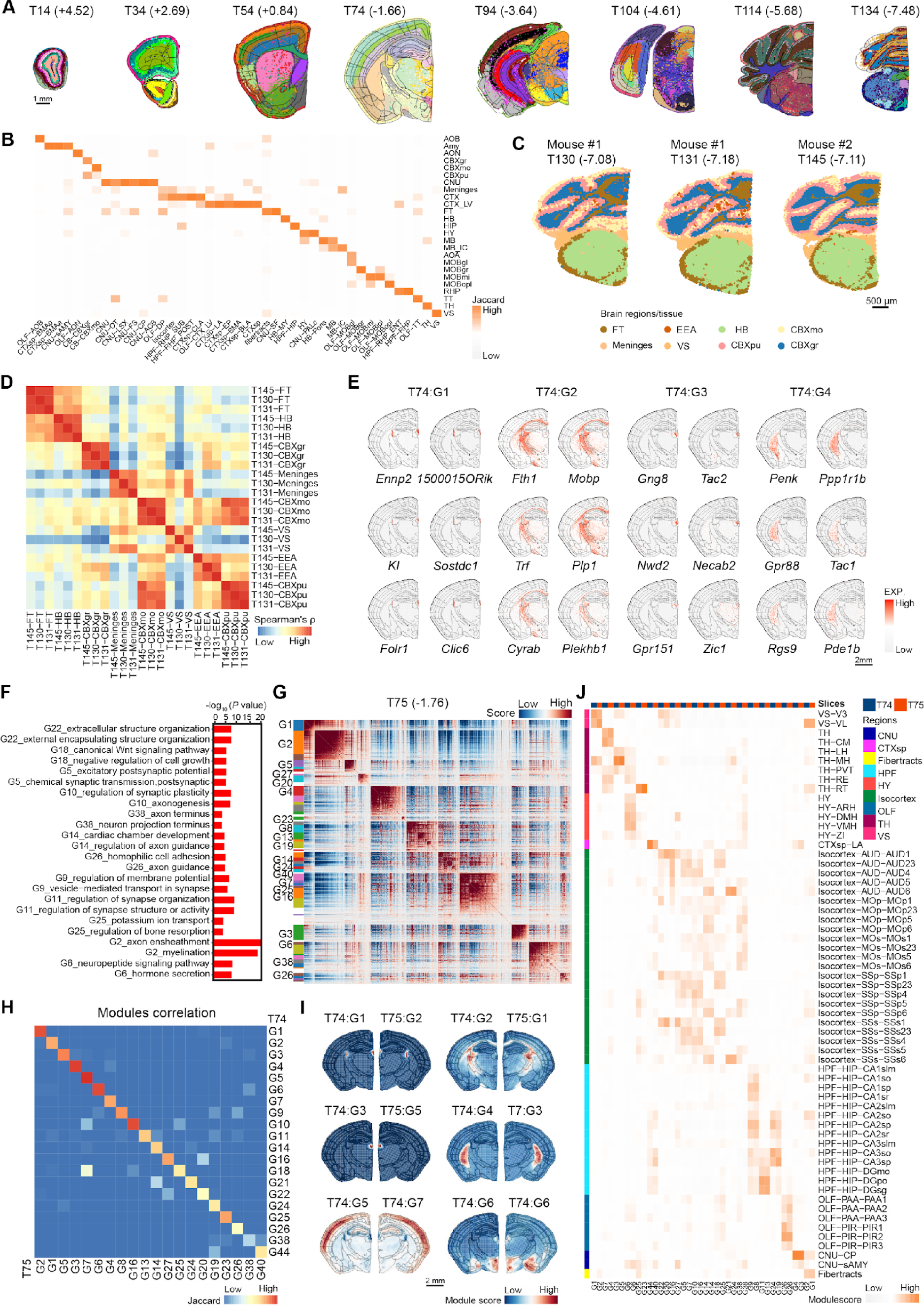
Reproducibility analysis of brain regions parcellation and reliability analysis of gene modules. **(A)**Spatial visualization of bin100 clusters in eight representative brain sections. Each color within the individual brain section represents a cluster generated by bin100 data. Scale bar, 1 mm. **(B)** Coincidence analysis of brain area parcellation in mice by clustering bin100 data compared with region boundaries based on ABA CCFv3. The horizontal axis represents the brain areas annotated after clustering of bin100 spots, while the vertical axis represents the areas of ABA CCFv3. **(C)** Spatial distribution of brain regions/tissue in three representative coronal brain sections from mouse #1 and mouse #2. Scale bar, 1 mm. **(D)** Spearman’s rank correlation coefficient (Spearman’s ρ) of brain regions partitioned across the three sections in Figure S5C. **(E)** Spatial visualization of top 3 genes from corresponding 4 gene modules in section T74. Scale bar, 2 mm. **(F)** Gene ontology functional enrichment analysis of brain region-specific gene modules in T74 brain sections. **(G)** Heatmap showing the genes with significant spatial autocorrelation grouped into different modules based on pairwise spatial correlations in brain section T75. **(H)** Heatmap showing the correlation between gene modules in T74 and T75 brain sections. Fisher’s exact test was performed between gene sets in each gene module. **(I)** Spatial visualization of the representative gene modules related to different brain regions in section T74 and T75. Scale bar, 2 mm. **(J)** Heatmap showing the spatial distribution of representative gene modules in the T74 and T75 brain sections.

**Figure S6.**
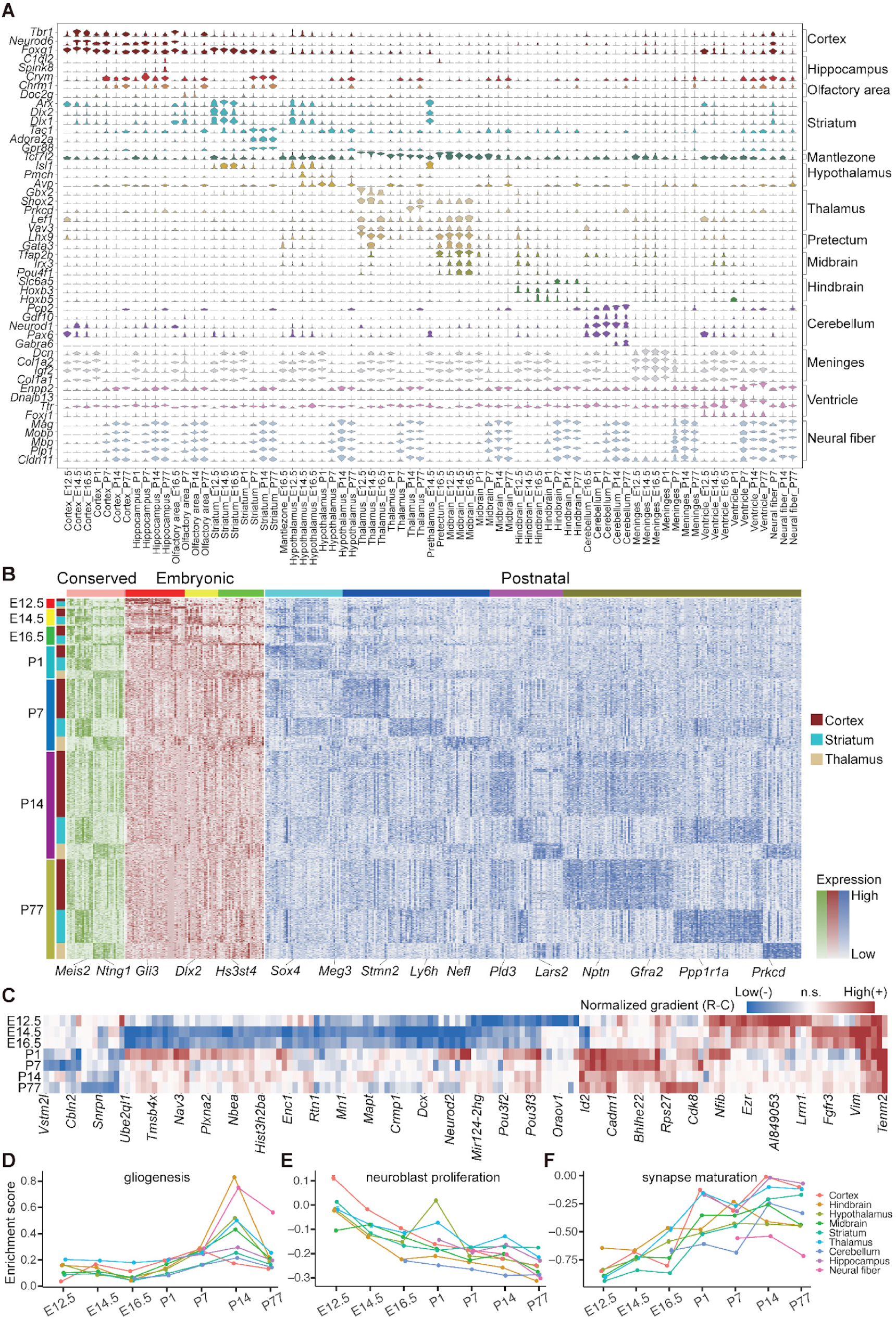
Spatiotemporal dynamics of gene expression in developing mouse brain. **(A)**Differentially expressed genes of brain region-specific clusters across developmental mouse brains (E12.5-P77). Violins are colored by their annotation. **(B)** Heatmap showing the expression of identified genes with conserved or time-dependent brain-regional specificity in developing brain regions (cortex, striatum, thalamus). **(C)** Heatmap showing the time-dependent expression gradient of indicated genes along cortical rostral-caudal axis in 7 developmental stages (E12.5-P77). Positive, negative gradients and non-significant ones are demonstrated with red, blue and white, respectively. **(D-F)** Gene enrichment score of neural development related GO biological processes including gliogenesis **(D)**, neuroblast proliferation **(E)**, and synapse maturation **(F)** across 7 developmental time points in 9 brain regions.

**Figure S7.**
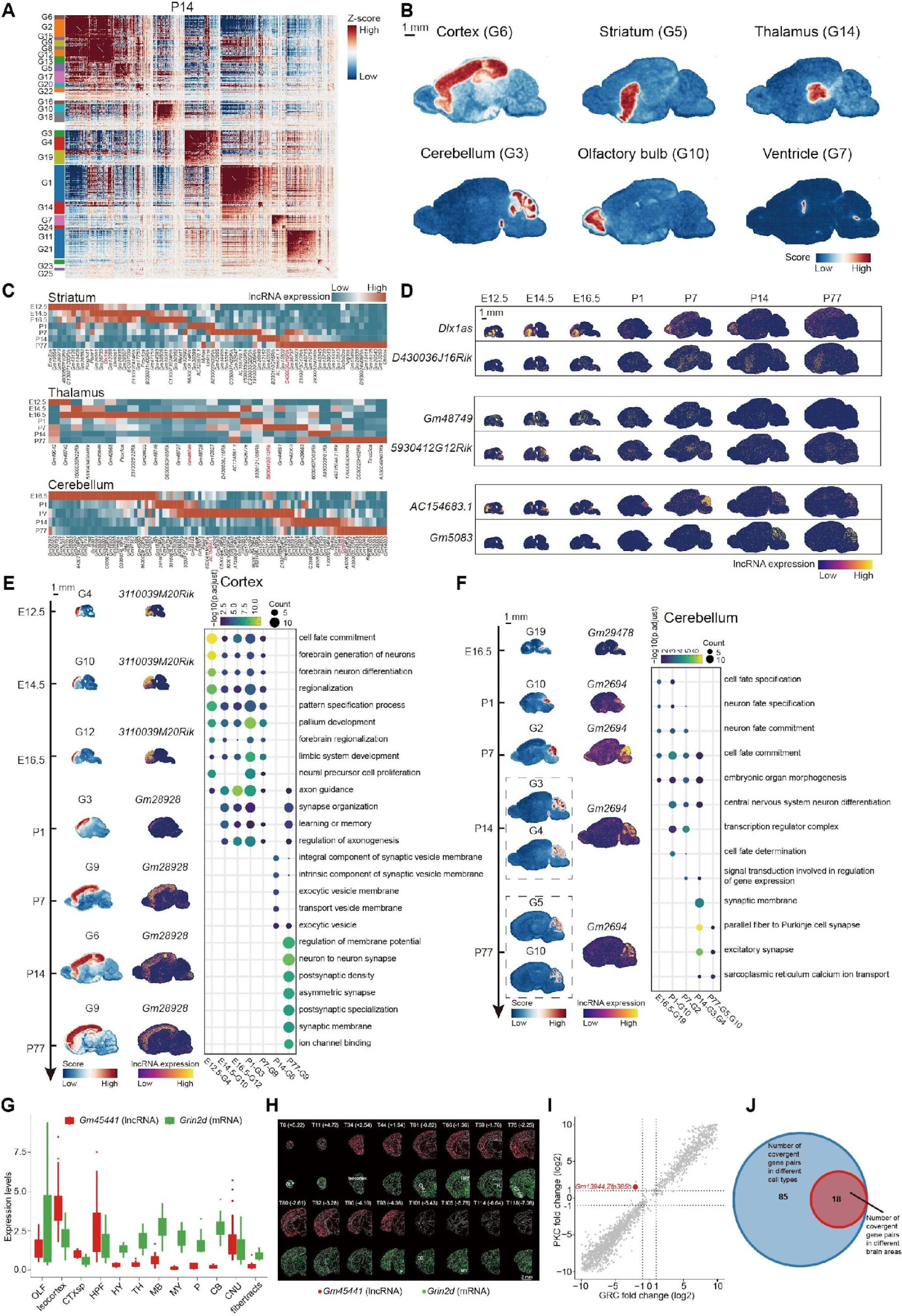
Spatial expression profile of specific non-coding RNA in the developmental mouse brain. **(A)**Heatmap of the genes with significant spatial autocorrelation grouped into different gene modules based on pairwise spatial correlations in P14 sagittal brain section. **(B)** Spatial distribution of 6 region-specific gene modules in P14 sagittal brain section. Scale bar, 1 mm. **(C)** Heatmap of the dynamic expression change of lncRNAs in developmental striatum, thalamus and cerebellum. **(D)** Spatial expression distribution of spatiotemporal brain region-specific lncRNAs (striatum, thalamus and cerebellum) at different developmental stages. Scale bar, 1 mm. **(E)** Left, Spatial distribution of cortex-specific modules and the expression of their corresponding lncRNAs (*3110039M20Rik, Gm28928*) at different developmental stages. Right, Bubble plot of the representative Gene Ontology enrichment pathways of cortex-specific modules at different developmental stages. Scale bar, 1 mm. **(F)** Left, Spatial distribution of cerebellum-specific modules and the expression of their corresponding lncRNAs (*Gm29478*, *Gm2694*) at different developmental stages. Right, Bubble plot of the representative Gene Ontology enrichment pathways of cerebellum-specific modules at different developmental stages. Scale bar, 1 mm. **(G)** Boxplot of the expression levels of the *Gm45441*-*Grin2d* convergent gene pair in different brain regions. **(H)** The spatial expression distribution in bin50 of *Gm45441-Grin2d* convergent gene pair. Section No. and bregma coordinates shown, unit mm. *Gm45441* is colored in red and *Grin2d* in green. Scale bar, 2mm. **(I)** The plot of fold changes of *Gm13944*-*Zfp385b* convergent gene pair between PKC and GRC neurons of cerebellum. **(J)** The overlap between brain area convergent gene pairs and cell type convergent gene pairs.

## Supplemental table titles and legends

**Table S1. Coronal sections information.**

**Table S2. Cell clusters annotation.**

**Table S3. Distribution of cell clusters ratio composition across brain areas.**

**Table S4. Classification of regional-specific genes in brainstem.**

**Table S5. Representative genes in spatial modules.**

**Table S6. Spatiotemporal profile of gene expression in the developing mouse brains.** (1) Spatiotemporal specificity of TF regulons in developing mouse brain. (2) Spatiotemporal specificity of genes in developing cortex, striatum and thalamus. (3) Temporal dynamics of TF regulons in developing cortex, striatum, thalamus and cerebellum. (4) Laminar enrichment and rostral-caudal gradient of TF regulons and genes in developing cortex.

**Table S7. The representative lncRNA in different brain regions and spatial modules.** Representative lncRNA with region-specific in different brain regions. (2) Representative lncRNA in spatial modules.

**Table S8. Gene modules enriched in brain regions during the developing mouse brains.**

**Table S9. The correlation of divergence and convergence lncRNA-mRNA gene pairs across all spatial transcriptome samples.**

**Table S10. Abbreviations of mouse brain areas.**

## Supplemental Movie titles

**Movie S1. 3D demonstration of selected cell clusters.**

**Movie S2. 3D demonstration of brainstem-specific neurons.**

**Movie S3. 3D demonstration of brainstem-specific neuropeptides.**

**Movie S4. 3D demonstration of region-specific lncRNAs.**

